# Joint recording of past and present transcriptomes reconstructs single-cell trajectories

**DOI:** 10.64898/2026.09.28.755048

**Authors:** Zeynep Aydin, Svetlana Ovchinnikova, Iris A Unterweger, Santiago Cerrizuela, Naomie Pont, Valentin Wüst, Ruben Boers, Katharina Zirngibl, Özgür Can, Matteo Spatuzzi, Tatiana Traboulsi, Xiaoyu Sun, Aylin Korkmaz, Sara Ortica, Tanya Foley, Katrin Volk, Susanne Kleber, Andres Sanz-Morejón, Milena Hasan, Joost Gribnau, Simon Anders, Ana Martin-Villalba, Laure Bally-Cuif

## Abstract

Tracing molecular cell state transitions through development, homeostasis and disease is still beyond reach at single-cell resolution. We present scReMeMber-Seq, a methylation-based cell-memory recorder connecting past transcriptional states with present transcriptome and DNA methylome in individual cells. scReMeMber-Seq leverages pulse induction of DCM-Polr2b (TimeMachine) and single-cell methylome-transcriptome (scMT) sequencing for endpoint readout. In mouse neural stem cells (NSCs) and pancreatic organoids, single-cell DCM methylation profiles (scDCMome) accurately discriminate cell types. Over time, scDCMome also record NSC quiescence-activation state changes across multiple cell divisions. In zebrafish embryos, scDCMome resolve individual transcriptional trajectories from progenitor to neuron. Together, scReMeMber-Seq enables real time reconstruction of cellular trajectories, connecting past and present molecular states in single cells, at genome scale, across experimental systems, *in vitro* and *in vivo*.

## INTRODUCTION

Biological systems are dynamic with cells transitioning through molecular states during development, immune responses, tissue homeostasis and repair. Derailments in these trajectories cause disease, while rerouting cell states offers therapeutic potential. Tracking these single-cell molecular trajectories is therefore essential to identify state transitions and their regulatory mechanisms. For this, one needs to link a cell’s past transcriptional activity to its present molecular identity.

One way to do this *in silico* is to reconstruct trajectories from single-cell transcriptomes spanning different stages of a transition, either from a steady-state sample that already contains such heterogeneity or by integrating data from multiple time points. Trajectory inference methods such as Slingshot (*1*), Destiny (*2*) or Monocle (*3*) find paths through the data’s latent-space representation that connect cells with similar transcriptomes, yielding a pseudotemporal ordering of cells, while approaches such as RNA velocity (*4*) and its variants (reviewed by (*5*)) add information on the direction of change. These methods, however, are inherently population-based: they cannot resolve the actual timing or speed of a transition, and they assume transitions to be gradual, requiring the full continuum of intermediate states to be represented by enough cells in the sample. Many biological processes instead involve fast or even abrupt changes, for example, daughter cells diverging sharply from their mother cell in an asymmetric division, that such computational inference cannot readily capture. This motivates methods that use direct approaches to actively follow cells’ lineage progression.

Several recent methods integrate temporal or lineage recording into single cells with endpoint transcriptomic profiling. They include longitudinal antibody-based cell labeling (*6*), CRISPR-Cas9-induced genetic scars resolved at single-cell (*7–9*) or tissue levels (*10*), and prime-editing-based sequential molecular recorders (*11*). The latter include signal-gated variants such as ENGRAM, which writes barcodes in bulk DNA when specified enhancers or signaling pathways drive pegRNA expression (*12*). These methods resolve lineage, temporal divergence and, when signal-gated, regulatory activity, but their record is limited to pre-designed molecular or temporal inputs rather than unbiased. None reconstruct a cell’s past transcriptome at single-cell resolution. Thus, gene-expression history remains unknown, and individual trajectories must instead be inferred computationally, leaving how cells transition between states unrecorded.

To date, single-cell transcriptome recording has been demonstrated in prokaryotes by converting selected RNAs into guide RNAs that write to DNA (*13*). In mammalian cells in culture, longitudinal transcriptomic profiling was successfully implemented by sampling small cytoplasmic volumes over time followed by Smart-Seq2 sRNAseq (*14*), but this method is not applicable *in vivo* nor in 3D culture models. Interesting methods based on the metabolic labeling of nascent RNAs with 4-thiouridine (4sU), such as scSLAM-seq or sci-fate allow reading old vs new RNAs in single cells, including in vivo (*15, 16*), but the temporal window is short and set by the stability of pre-existing transcripts. Finally, recently, transcriptome-wide RNA storage in mammalian cells during a defined recording window has become possible with TimeVault (*17*); however, it lacks joint readout of past transcriptional history and present transcriptomic/epigenomic states in single cells. HisTrac-seq narrows this gap by using inducible Dam to bookmark past transcription as DNA methylation, later co-reading it with current H3K27ac via dual CUT&Tag in single cells. Yet this joint readout has only been shown in vitro; in vivo, only past-only bookmarking was demonstrated (*18*). Consequently, key priming programs often remain unobserved, weakening causal links between what a cell {was and what it is now. Here we present scReMeMber-Seq, a molecular cell-history recorder that reconstructs past transcriptional states while jointly profiling present transcriptomic and methylomic landscapes at single cell resolution. scReMeMber-Seq leverages the TimeMachine principle described by Boers et al., where E. coli DNA cytosine methyltrans-ferase (DCM) fused to the RNA polymerase II subunit methylates CmeC(A/T)GG sites at actively transcribed loci (*19*). Integrated with an induction system and single-cell Methylome and Transcriptome (scMT) sequencing (*20*), the method enables pulse–chase, lineage-specific and/or temporal labeling of past transcriptional activity together with present-state profiling. Applying scReMeMber-Seq to adult mouse lineages in culture, we identified pancreatic and neural cell types solely from DCM methylation marks. We also captured current neural stem cell (NSC) states and determined whether individual cells had been active or quiescent in the past. Applied to zebrafish embryonic neurogenesis *in vivo*, DCM signatures could identify the neural lineage against all other cell types and connect individual neurons to their progenitor state of origin. Finally, in both systems, reading the present cells’ state includes single-cell methylome information.

## RESULTS

### Finding conditions for best signal-to-noise ratio of DCM induction in mouse neural stem cells *in vitro*

The scReMeMber-Seq approach is based on single cell multi-omics profiling of cells expressing an inducible fusion of the bacterial DNA cytosine methyltransferase DCM to Polr2B (*19*) (Fig. 1). We first checked that the zebrafish and mouse genomes are suitable for the recording of DCM marks in single cells (Supplementary text and Fig. S1). Thereafter, we tested scReMeMber-Seq on a culture of actively cycling murine neural stem cells (NSCs) isolated from a Doxycycline (Dox)-inducible DCM-Pol2b fusion mouse line (*19*). Profiling was achieved with scMT-seq (see Supplementary text). We found that only few DCM motifs were methylated at baseline, and Dox-induced DCM expression noticeably increased the genome-wide fraction of methylated DCM motifs (“global methylation fraction”) (Fig. S2A).

**Fig. 1.**
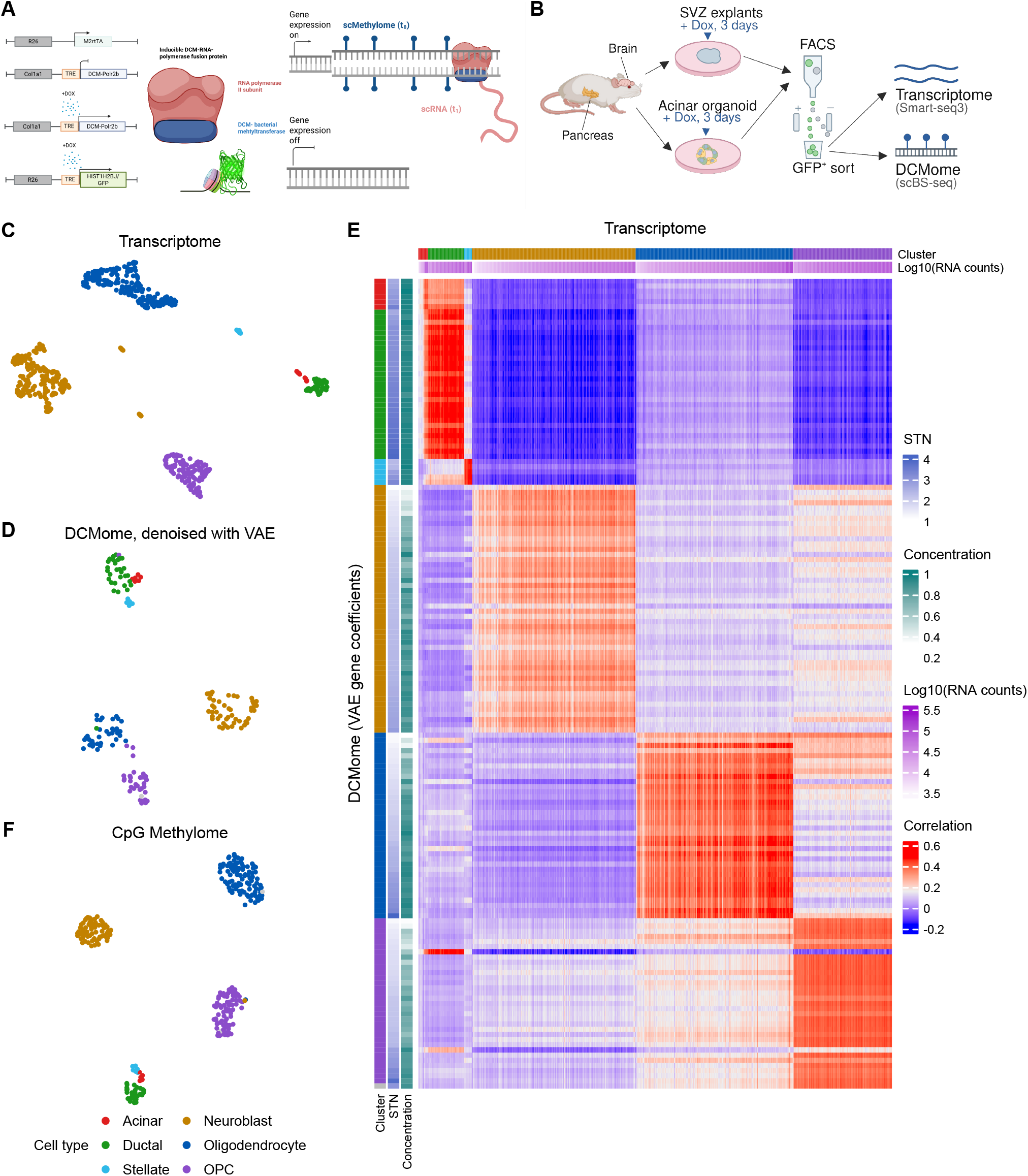
The DCMome alone enables distinction of cell types. **(A)** Principle of scReMeMber-Seq: Inducible expression of DCM (blue) fused to the RNA Polymerase IIb subunit (red) leads to methylation of DCM motifs in actively transcribed genes. **(B)** Schema of the explant and organoid experiment demonstrating cell-type identification from scDCM marks. **(C)** UMAP representation of the latent space derived from the cells’ transcriptomes with PCA. Colors mark Louvain clusters, labels show cell types assigned to the clusters. **(D)** UMAP representation of the latent space derived from the VAE-denoised cells’ DCMome with PCA. Colors as in (C). **(E)** Heatmap of correlation coefficients between each cell’s transcriptome (columns) and every cell’s VAE-denoised DCMome (rows). Side bars show, for transcriptomes: read number and cell-type cluster; for DCMomes: cluster, signal-to-noise ratio, and Dirichlet concentration parameter (a quantity indicating signal quality as perceived by the VAE). **(F)** UMAP representation of the PCA embedding derived from the endogenous DNA methylation (CpG methylation), produced using the MethSCAn (*24*) pipeline.

**Fig. 2.**
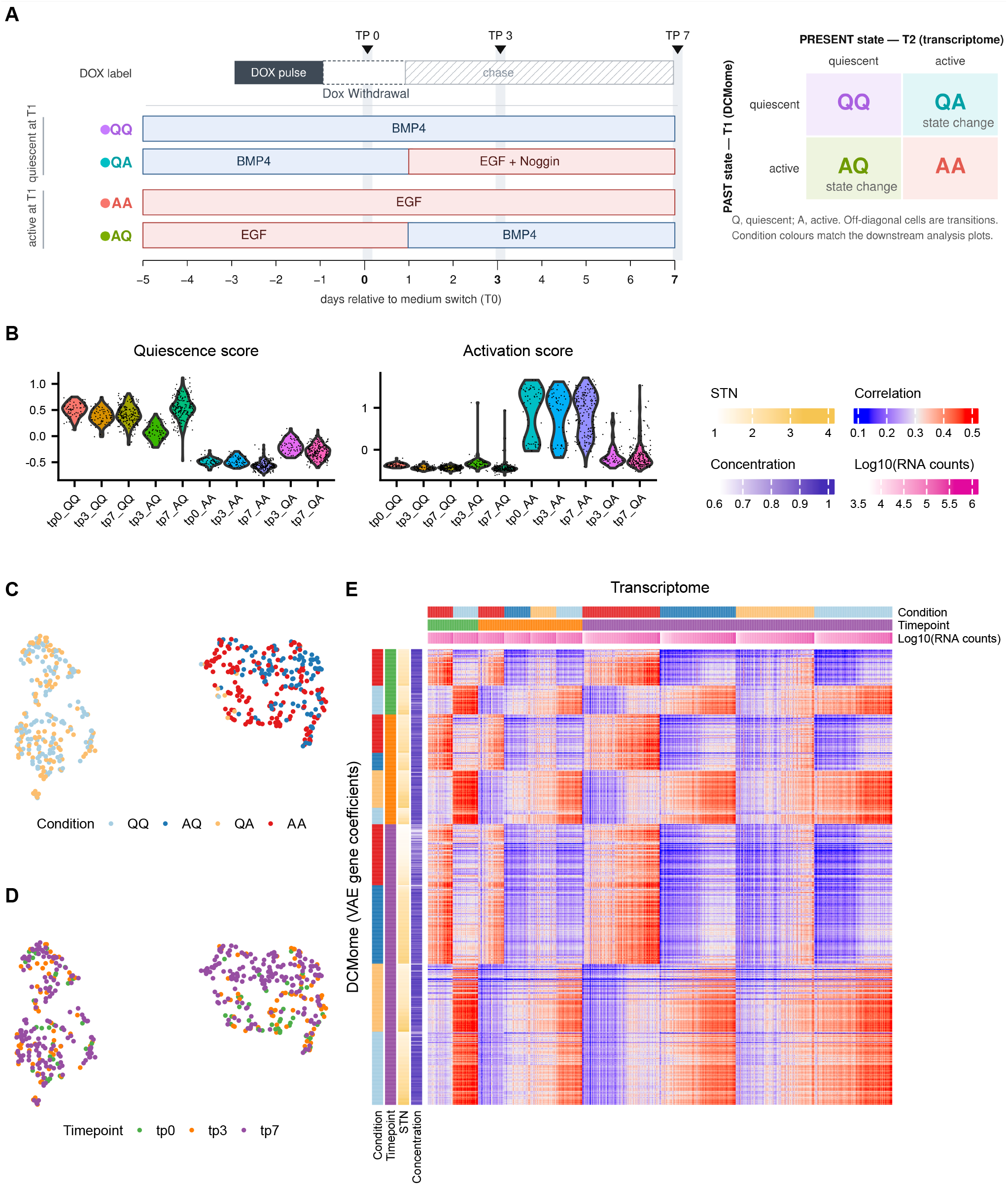
scReMeMber-Seq enables identification of state of origin of neural stem cells in vitro. **(A)** Schema of the pulse-chase time course with condition switch. **(B)** Beeswarm plots showing A (activation) and Q (quiescence) score for the cells, stratified by samples (i.e., by experimental condition and time point). **(C,D)** UMAP representation of the VAE-derived latent embedding of the cells’ scDCMomes, coloured by experimental condition (C) and by sampling time point (i.e., length of chase) (D). **(E)** Correlation heatmap comparing all cells’ transcriptomes with the cells’ scDCMomes.

**Fig. 3.**
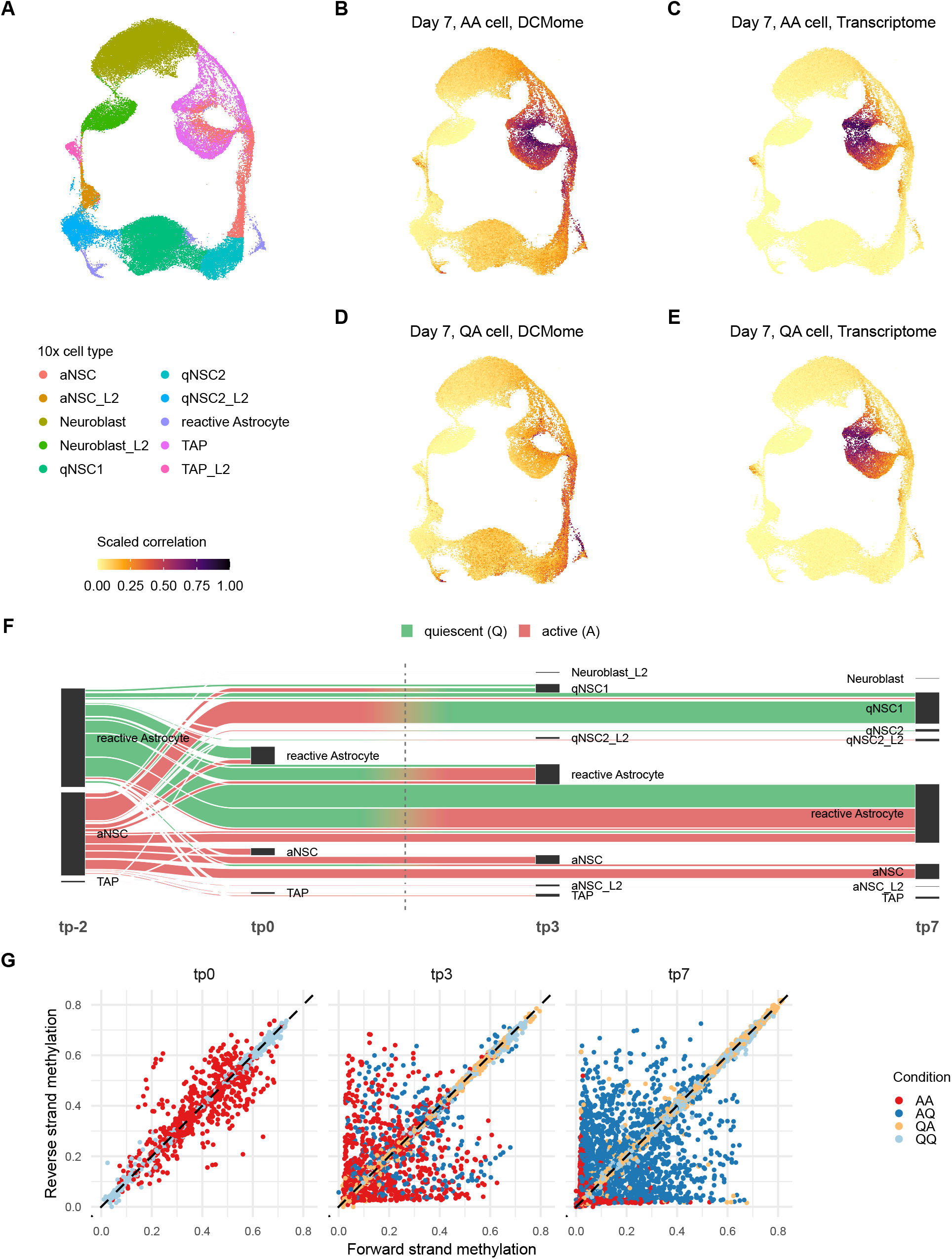
scReMeMber-Seq enables reconstruction of state transitions of neural stem cells in vitro. **(A)** Atlas of murine subventricular zone cells (astrocytes, NSCs and their GLAST-expressing progeny), from 10X droplet scRNA-seq, visualized as UMAP. **(B,C)** Correlation of one TP7 AA cell’s scDCMome (B) and transcriptome (C) with all atlas cells; color indicates correlation strength. **(D,E)** Same as (B,C) but for a TP7 QA cell. **(F)** Sankey plot illustrating lineage progression, linking the cell identity inferred from the scDCMome (left) to the transcriptomic identity at the time of collection (to the right). See also interactive Sankey plot, stratified by sample condition and time point, at this link. **(G)** Comparison of methylation of forward and reverse strand. Each point is one chromosome (autosome) of one cell, i.e. each cell is represented by several points. Colours indicate experimental condition; time points are given in the plot titles.

**Fig. 4.**
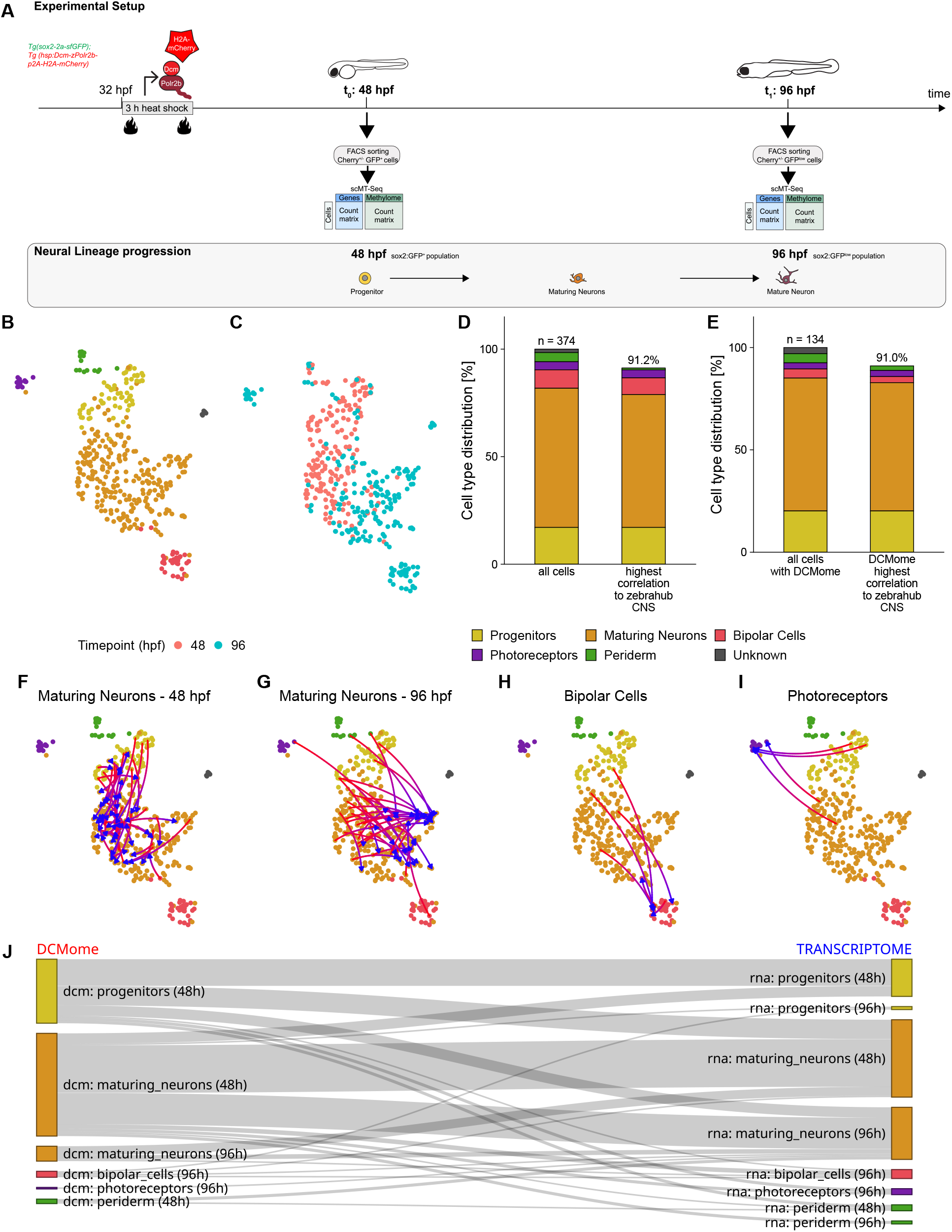
scReMeMberSeq enables reconstruction of the developmental neural lineage *in vivo* in zebrafish. **(A)** Schematic overview of the experimental setup. scReMeMberSeq was applied to the neural lineage during zebrafish embryogenesis using a pulse-chase strategy. DCM labeling was induced by repeated heat shock at 32 hours post-fertilization (hpf). Single cells were collected at 48 hpf or 96 hpf for scReMeMberSeq. **(B)** Transcriptomic UMAP of sorted single cells of 48 hpf and 96 hpf, colored by terminal cell type identity. **(C)** Transcriptomic UMAP colored by the time of collection, either 48 hpf (red) or 96 hpf (blue). **(D)** Bar chart showing the cell type distribution of all sorted cells (n = 374) and the proportion of cells whose transcriptome shows the highest correlation to a CNS cell in the Zebrahub reference dataset). **(E)** Bar chart showing the cell type distribution of all sorted cells with a DCMome (n = 136) and the proportion of cells whose transcriptome shows the highest correlation to a CNS cell in the Zebrahub reference dataset. **(F-I)** UMAP in which cells are color-coded according to their cell identity (B). Arrows connect the transcriptomic position of a given cell (blue arrowhead) to the cell in the UMAP whose transcriptome most closely correlates with that cell’s DCMome (red arrowbase). Arrows are shown for: maturing neurons at 48 hpf (F), maturing neurons at 96 hpf (G), bipolar cells (H), and photoreceptors (I). **(J)** Sankey plot illustrating lineage progression, linking the cell identity inferred from the scDCMome (left) to the transcriptomic identity at the time of collection (right).

We quantified specificity as a “signal-to-noise ratio” (STN): the ratio of the methylation fraction inside genes vs in intergenic regions (motifs at least 5 kb from any annotated gene or enhancer). As expected, uninduced Polr2B-DCM cells had an STN near 1, and Dox induction increased both global methylation and STN (Fig. S2B) until methylation reached 15-20% of motifs.

Testing a range of Dox concentrations and pulse durations, we found pulse length, not dosage, drove both metrics (Fig. S2C,D): only 48h pulses reliably gave clear gene-specific methylation (STN>1.5). Chase experiments showed methylation continued accumulating for 24h after pulse termination and began to decline again by 48h (Fig. S2E). This matched the expression of DCM-Polr2B protein, which peaked at 48h Dox treatment and declined thereafter (Fig. S2F).

### Construction and validation of an in-vivo inducible DCM-Polr2b methylation marking system in zebrafish

To apply scReMeMber-Seq in zebrafish, we built a transgenic line expressing a fusion of DCM with zebrafish RNA Polymerase II subunit B (Polr2b) under control of the zebrafish *heat-shock 70* promoter (*21*). We added H2A-mCherry to stably track cells having expressed the transgene (Tg(*hsp:DCM-polr2b-P2A-H2A-mCherryip19*)) (Fig. S3A). To focus on the neuronal lineage, we crossed this line into Tg(*sox2-2a-sfGFPstl8*) (*22*), labeling neural progenitors (Sox2pos) and tracking their progeny over several days via sfGFP protein stability. To ensure homogeneous transgene expression levels between embryos, we worked with heterozygous embryos for both backgrounds. Upon heat-shock (hs), all tissues visually induced the transgene (Fig. S3B,C), and mCherry persisted over time (Fig. S3D). FACS of all cells from non-heat-shocked vs heat-shocked embryos also showed a quasi-absence of leakiness, as there were only very few H2A-mCherry-positive cells without hs (Fig. S3E-G). Finally, we assessed the dynamics of transgene induction using two 45-minute hs pulses within 2 hours starting in embryos at 30 hours-post-fertilization (hpf), followed by time-course quantification of *DCM* transcripts by in situ hybridization chain reaction (HCR). Shortly after the hs pulse, at 34hpf, high *DCM* expression levels were already detected in most embryonic cells (Fig. S3H,I). Transcript levels then gradually decreased back to baseline by 48hpf, comparable between wildtype and non-induced embryos (Fig. S3J).

Next, we tested whether a hs pulse was sufficient to induce specific DCM methylation in the genome. We maximized transgene induction over a short time frame with two 45-minute hs pulses at 90 min interval starting at 32hpf, then collected embryos at 48hpf. Control embryos were wildtype embryos that were also heat-shocked. Individual H2A-mCherrypos,GFPpos cells were sorted by FACS and processed for scMT-seq (*20*). A few minibulks of 25 cells each were processed in parallel. The global DCM methylation fraction was increased in individual cells and minibulks from heat-shocked transgenic compared to wildtype embryos (Fig. S3K). DCM methylation levels correlated with H2A-mCherry intensity at the cell collection timepoint (Fig. S3L). Finally, the DCMome STN of most cells was higher upon induction than in control cells, confirming that DCM methylation was preferentially driven to gene bodies by the DCM-Polr2b fusion (Fig. S3M).

### Upon induction of DCM-Polr2b, increased methylation at gene bodies is detectable at the level of individual genes in single cells

In zebrafish, DCM methylation was enriched at genes expected to be expressed in the neural lineage at 48 hpf (Fig. S4A-B). Likewise, in mouse, enrichment was observed at tissue-specific marker genes actively expressed in the induced tissues (Fig. S4C). Comparing the pooled transcriptomes of minibulks from heat-shocked embryos at 48hpf with vs without DCM-Polr2b construct, we also found that DCM induction and methylation had no global effect on gene expression in zebrafish (Fig. S4D), as previously shown for mouse (*19*).

### The scDCMome reliably identifies cell types in mouse

To test whether the scDCMome alone resolves cell types, we benchmarked within- and between-tissue divergence using brain and pancreas, where cell types are related within but distinct across tissues. Subventricular zone (SVZ) explants and acinar-derived organoids (ADO) from the same mouse were labeled in parallel with 8 µg/mL Dox for three days, and GFPhigh cells were FACS-sorted and processed for scMT-seq (1B). We first performed a standard transcriptome-based analysis: log-normalized mRNA counts were PCA-transformed and visualized as a Louvain-clustered UMAP (1C). We identified the expected diversity in each preparation: acinar, ductal-like, and stellate cells in ADOs, and neuroblasts, oligodendrocytes, and oligodendrocyte progenitors in SVZ explants.

To determine whether the scDCMome alone could yield similar cell-type separation, we used a variational autoencoder (VAE) instead of PCA for the latent representation. The resulting UMAP (1D) shows clear separation of the transcriptome-defined cell types; thus, the scDCMome carries sufficient information for cell-type identification. 112 cells with weak signal were excluded, among which 78 were excluded based on the low promoter methylation (Fig. S5A,B). As a lighter-weight alternative to the VAE, we also used PCA applied to the binomial deviance of each gene’s DCM methylation from an estimated background. This performed well for high-signal cells but underperformed the VAE for weak signals (Fig. S5C,D). To test whether scDCMome differences actually reflect transcriptomic differences between cell types for the same genes, we correlated every cell’s transcriptome with every cell’s DCMome (1E): scDCMome and transcriptome were most similar within, not between, cell types. These correlations were stronger using VAE-denoised DCMome values than “raw” deviances (Fig. S5E), consistent with the ability of VAEs to denoise data. Together, scReMeMber-Seq enables robust cell-type discrimination from DCMome alone, without gene expression information.

Finally, scMT-seq also profiles the endogenous DNA methylome at CpG sites, which likewise separates cell types (1F).

### scReMeMber-Seq enables tracing transcriptome history across state transitions and cell cycles in individual mouse NSCs

We next asked whether a cell’s prior state, as recorded in the DCMome, can be reconstructed after a chase, and for how many cell divisions it remains readable. NSCs were cultured in media keeping them either actively cycling (A) or quiescent (Q), then either maintained or switched to the other condition (2A), giving four trajectories: AA (cells remain active and cycling), QQ (cells remain quiescent), AQ (cells transit from active to quiescent), and QA (cells transit from quiescent to active). We assessed each cell’s DCM-recorded initial (“past”) condition alongside its transcriptional state at harvest (“present”) at the indicated time points (2A). We first assessed the current state as indicated by the transcriptome, using a “Q score” and an “A score” (averaged expression of gene sets upregulated by BMP or EGF exposure, respectively) as in an earlier NSC experiment (*23*). QQ and AA cells scored as quiescent and active as expected, as did AQ cells (quiescent); QA cells appeared in transition, with Q score already low but A score not yet fully risen (2B). We then examined the scDCMome, again using a VAE-denoised latent embedding. Cells clustered by initial condition but mixed by final condition (2C), showing that the scDCMome reflects the Dox-pulse condition. Good mixing across sampling time points confirmed the record’s stability after washout (2D). A correlation heatmap confirmed that scDCMomes were always more similar to transcriptomes matching their pulse condition (2E).

### Comparison with an atlas reveals fine-grained cell-state annotation in past and present

To test whether scReMeMber-Seq resolves finer cell-state differences, we generated a 10X droplet scRNA-seq atlas of the SVZ (3A) and correlated each culture cell’s DCMome or transcriptome with the transcriptome of every cell in the atlas. The scDCMome, and the transcriptome, of active cells was highly correlated with actively cycling transient amplifying progenitors and neuroblasts (3B,C), whereas for some QA cells the sc DCMome was spread between quiescent state (qNSC1) and reactive astrocytes, and the transcriptome correlated with highly cycling cells (3D,E). Assigning each culture cell thus to its best-correlated atlas cell type let us cross-tabulate these transitions as a Sankey plot (3F; link to interactive version); an alternative color-gradient view of individual mappings is shown in Fig. S6A-E), linking past and present, and tracing state transitions in real time. Notably, in conditions with no transition induced, the transcriptome and the DCMome matched each other, while otherwise transitions could be tracked by discordant reasdouts. BMP4 exposure locked most cells into a reactive state, while a few entered the qNSC1 state. Subsequent exposure to activating medium (EGF/Noggin), allowed a small subset to progress to shallow quiescence (qNSC2) and to aNSCs. Similarly, we previously reported that following brain ischemia, only a fraction of reactive astrocytes enters the qNSC2 state and acquires neurogenic competence (*24*). Interestingly, a shorter treatment with BMP4 forces active cells into quiescence rather than to a reactive state. Together, these results show that scRe-MeMber-Seq can resolve trajectories in real time, even in actively dividing cells for up to 7 divisions (considering a cell cycle of 17.5h (*25*) and see below 3G).

### The VAE-based analysis can compensate for signal loss due to cell divisions

The VAE was essential for achieving separation by past state in 2C,D; the equivalent binomial-deviance-based plots look markedly worse (Fig. S6F-H). This is in part due to how the DCM signal dilutes with divisions: both DNA strands are methylated during the pulse, but strands synthesized afterward are created with no marks. Boers et al. (*19*) have observed that marks can be copied during DNA duplication, but the possible perdurance of Dcm activity during the first cycle(s) in our study is difficult to assess. Thus, each division halves (or reduces) the number of marked chromatids on average, and (barring recombination) any given strand remains either fully marked or significantly less marked. Accordingly, overall DCM methylation correlated with the number of cell cycles, assessed by dilution of GFP-H2B (Fig. S7A-C). DCM methylation was lower in cells that cycled during the chase (Fig. S7D), with strongest reduction in AA, followed by AQ (incomplete transition to quiescence) then QA (few cycles, activation not being complete).

This is clearer per chromosome strand (3G): at TP0, quiescent cells show balanced marks on both strands, while active cells that divided during Dox exposure are imbalanced (post-division marks cover both strands, earlier marks only one). After further chase, non-dividing cells (QQ, and most QA) stay balanced (diagonal in 3G), while cells dividing during the pulse (QA, AA) become imbalanced. Divisions after Dox activity ceased can leave one fully empty strand, seen as points on the axes for AA at TP7 and, partially, TP3.

We exploit this by computing each strand’s methylation fraction relative to other strands as a signal-content measure. For PCA/binomial-deviance, strands well below the cell’s best strand are excluded; for the VAE, strand signal content enters the loss function so the autoencoder is not penalized for dilution-related missing marks, encouraging it to impute them by borrowing from similar cells’ well-marked strands. In summary, the VAE provides both denoising and implicit imputation of information lost to cell division, greatly extending our method’s usability: DCM profiles remain identifiable even when divisions have erased signal from all but a few chromosome strands.

### scReMeMber-Seq enables tracing transcriptome history *in vivo* during embryonic neurogenesis in zebrafish

To test whether scReMeMber-Seq can also distinguish different cell states in an *in vivo* model with rapid temporal dynamics, we studied neural lineage progression during zebrafish embryonic development. We induced DCM methylation by heat shocking *Tg(hsp:DCM-polr2b-P2A-H2A-mCherryip199 sox2-2a-sfGFPstl8*) embryos with two pulses starting at 32hpf. Individual H2A-mCherrypos,GFPpos cells were FACS-sorted and processed for scMT-seq at both 48hpf and 96hpf (4A). We chose 48hpf (t0) to capture cells at maximal DCM labeling following hs induction, a time point when Sox2pos neural progenitor cells reside predominantly in an undifferentiated progenitor state (36-48hpf). By 96hpf (t1), cells were expected to have differentiated into more mature neuronal fates through no more than 2–3 cell divisions, during a period in which no *de novo* DCM labeling would occur. To further enrich for mature neurons, we FACS sorted specifically GFPlow cells at 96hpf (Fig. S8A,B).

The transcriptomes of all sampled cells revealed a continuum of differentiation states along the neural lineage (4B). Most cells at 48hpf were neural progenitors or cells just beginning to express early differentiation markers, such as *elavl3/4*, whereas at 96hpf the majority of cells were maturing neurons and mature neuronal cell types, including photoreceptors and bipolar cells (4B,C). As Sox2pos cells are not synchronized at the time of labeling, overlapping cell states were observed at both time points in different proportions (4C, Fig. S8C) and within each cluster. As clusters 0 and 1 showed extensive overlap, we merged them and designated the combined cluster ‘maturing neurons’ (4B); 48 and 96hpf cells within this merged cluster occupied partially distinct positions, reflecting progressive maturation (4C). Additionally, we identified a small cluster of Sox2pos periderm cells and a small cluster of contaminating cells, containing a mixture of microglia (*mpeg* pos) and hematopoietic cells, which we labelled as unknown and excluded from further analysis (4B). Individual clusters showed high transcriptomic overlap, with only a small proportion of genes uniquely expressed within a single cluster, consistent with the continuous nature of neural differentiation (Fig. S8D). Both control and DCM-labeled cells were present across all clusters, demonstrating that DCM expression itself did not induce a distinct cell state (Fig. S8E).

DCM methylation levels showed a weaker correlation with H2A-mCherry protein intensity at 96hpf compared to 48hpf (Fig. S8F, compare with Fig. S3L), most likely reflecting dilution of the DCM label through successive cell divisions. Consistently, the STN at 96hpf was reduced, with only a subset of cells showing values above those of control cells (Fig. S8G,H, compare with Fig. S3M).

To determine whether the scDCMome could predict cell type, we developed a quantifiable cell type assessment. We first calculated correlations between transcriptomic signatures in our sampled cells with the Zebrahub atlas (*26*), containing cells ranging from end-of-gastrulation whole embryos to 10-day old larvae. Each of our cells, besides those in the ‘periderm’ and ‘unknown’ clusters, showed highest transcriptome correlation to a cell categorized as ‘central nervous system’ between 1 and 5dpf in the Zebrahub atlas (4D). Next, we addressed the reliability of the scDCMome alone to similarly predict cell type, by substituting the scDCMome for the transcriptome in otherwise identical correlation calculations. 90% of cells had highest DCMome correlation to the transcriptome of a cell from the central nervous system in the Zebrahub atlas (4E, example in Fig. S9A-D, link to interactive viewer). Thus, the scDCMome specifically captures actively transcribed genes at the time of induction and is sufficient to predict overall cell type identity.

Finally, we assessed whether comparing scDCMome and transcriptome could reconstruct the *in vivo* transcriptional history of individual cells. For this, we correlated each cell’s DCMome to the transcriptomes of all profiled cells and visualized the results on the UMAP. The scDCMome of ‘maturing neurons’ collected at 48hpf corresponded most closely to ‘progenitor cell’ transcriptomes (4F), whereas the scDCMome of ‘maturing neurons’ collected at 96hpf matched most strongly to 48hpf ‘maturing neuron’ transcriptomes (4G), revealing a maturation trajectory within this cluster across time. The scDCMome of the most terminally differentiated cell types (bipolar cells and photoreceptors) mapped almost exclusively to ‘progenitor’ states (4H,I, Fig. S9E,F). Together, these results reveal a consistent temporal shift across the neural lineage, where most cells’ transcriptome reflects a more mature state than their scDCMome (4J and link to interactive version). as expected if DCM methylation captures gene activity at an earlier developmental stage. Taken together, this benchmark establishes scReMeMber-Seq as a tool capable of reconstructing transcriptional histories at single-cell resolution during *in vivo* neurogenesis.

## DISCUSSION

scReMeMber-Seq achieves genome-wide, unbiased transcriptional recording at single-cell resolution by exploiting inducible expression of DCM, which deposits transcriptional memory marks (the DCMome), read out together with the transcriptome and methylome of the same single cell. In contrast to current methods (*6, 9, 10, 14, 17, 18*), scReMeMber-Seq requires no prior target selection and jointly reads past and present in the same cell. Because this reflects real transcriptional history rather than an inferred one, it also needs no defined starting point, circumventing the directionality imposed by pseudotime methods, and allowing us to reconstruct prior cell states within an unsynchronized, progressively differentiating population *in vivo*.

Whole-genome methylation profiles are inherently sparse at single-cell resolution, a limitation further compounded by dilution of DCM marks through DNA replication in proliferating populations. We addressed this by exploiting strand segregation of DCM methylation per chromosome and analyzing the resulting signal with a VAE, which recovers usable signal even from cells with very low methylation coverage and correlates it with the corresponding transcriptome.

The joint readout of past and present states extends what single-cell methylome-transcriptome (scMT) sequencing has begun to reveal: cells with similar transcriptomes can carry distinct methylomes, as we previously showed for the astrocyte-to-quiescent-NSC transition (*24*). By coupling scMT-seq to DCM-Pol2b–based recording, scRe-MeMber-Seq adds the missing temporal axis, making it possible to ask, for the first time, which initial state determined a cell’s present state or fate. This should offer mechanistic insight into lineage commitment, plasticity, and reprogramming beyond static or trajectory-inferred snapshots, and generalize to other systems where cells change identity over time, including development, tissue regeneration, immune activation and exhaustion, and the gradual drift of cell states in aging and disease.

## Supporting information

Full supplementary material

## Acknowledgements

For data exchange, we used SDS@hd, a service funded by the State of Baden-Württemberg (Ministry of Science, Research and the Arts) and the DFG (grant INST 35/1503-1 FUGG. For computational resources the S.A. lab was supported by the de.NBI Cloud within the German Network for Bioinformatics Infrastructure (de.NBI) and ELIXIR-DE (Forschungszentrum Jülich and W-de.NBI-001, W-de.NBI-004, W-de.NBI-008, W-de.NBI-010, W-de.NBI-013, W-de.NBI-014, W-de.NBI-016, W-de.NBI-022). The A.M-V lab is grateful to the DKFZ genomics and proteomics core Facility; the DKFZ sequencing open laboratory, the DKFZ Single Cell Open Lab and the DKFZ Flow Cytometry facility, in particular Diana Ordonez Rueda. The L.B-C lab is grateful to Marc Monot (Biomics platform) and Sandrine Schmutz (Flow Cytometry platform) at Institut Pasteur for advice on sequencing and FACS, respectively, to all ZEN lab members for insight and support, and to Sébastien Bedu and Nathan Guibert for expert fish care and help with maintaining the zebrafish transgenic lines generated in this study.

## Funding

This work was funded by ERC (AdG SyG 595 101071786 – PEPS) (to S.A, L.B-C and A.M-V). Additional funded was provided by the Klaus Tschira Foundation (grant 00.022.2019) (to S.A), the DFG-RTG2727-445549683-In Check (to A.M-V), and the German Cancer Research Center (DKFZ) (to A.M-V), and the Agence Nationale de la Recherche (Labex Revive ANR-10-LABX-0073 and a government grant managed by the ANR under the France 2030 program with reference numbers ANR-24-EXCI-0001, ANR-24-EXCI-0002, ANR-24-EXCI-0003, ANR-24-EXCI-0004, ANR-24-EXCI-0005) (to L.B-C).

## Authors’ contributions

Experimental work : Z.A., Ö.C., S.C., T.F., A.K., S.K., A.S-M, S.Or., T.T., I.A.U., K.V.; Computational analysis: S.Ov., N.P., M.S., X.S., V.W.; Resources : R.B., J.G., M.H. ; Writing, review, editing: S.A., Z.A., L.B-C., A.M-V., S.Ov., N.P., I.A.U.; Conceptualization, funding acquisition: S.A., L.B-C., A.M-V. ; Supervision: S.A., L.B-C., A.M-V., A.S-M.

## Competing interests

The authors declare that they have no competing interests.

## Data, code and material’s availability

All source codes produced and used in the analysis are available at this link. The zebrafish transgenic line generated in this study is available upon request to L.B-C.

## MATERIAL AND METHODS

### Experimental work in mouse

#### Mouse models and animal welfare

All animal work was performed in accordance with institutional and governmental regulations and was authorized by the Regierungspräsidium Karlsruhe, Germany. Animals were housed in the animal facilities of the German Cancer Research Center (DKFZ) at a 12 h dark/ light cycle with free access to food andwater. Humidity was kept at 55% and temperature at 22 °C. The following mouse lines were used: C57BL/6N (for wild-type control) and Gt(ROSA)26Sor^tm1(rtTA*M2)Jae^ Col1a1^tm1(tetO-DCM-PolIIB)Grib^Tg(tetO-HIST1H2BJ/GFP)47Efu alleles (TetO-DCMpolIIB-H2BGFP line; for retrospective gene expression analysis). Mice were male and were age-matched to 2 months old.

#### Neural stem cell culture

NSCs were isolated from the ventricular–subventricular zone (SVZ) of 8–12-week-old male mice euthanized by cervical dislocation. Brains were transferred on ice into HBSS (Life Technologies) supplemented with 2.5 mM HEPES (ThermoFisher), 0.65% D-glucose, and 1% penicillin–streptomycin (Invitrogen). The vSVZ was dissected as described (Mirzadeh et al., 2010) and dissociated using papain (Neural Tissue Dissociation Kit, Miltenyi Biotec). Cells were maintained as neurospheres in Neurobasal A supplemented with 2% B27, 1% L-glutamine (ThermoFisher), 2 μg/mL heparin, 20 ng/mL bFGF (ReliaTech), and 20 ng/ mL EGF (Promokine), at 37 °C and 5% CO_2_. Spheres were dissociated with Accutase (ThermoFisher) after 5–7 days and passaged every 3–4 days.

#### Western blot and DCM protein degradation assays on mouse NSCs

DCM-Polr2b-positive NSCs, isolated from the subventricular zone (SVZ) at passage 10, were treated with Doxycycline (4 µg/mL, 48 h) and collected at 24 h and 48 h of treatment, then at 0, 1, 3, and 7 days post-withdrawal (Tp0, c1, c3, c7), alongside an untreated negative control. Whole-cell lysates (25 µg protein/sample) were resolved by SDS-PAGE, transferred to nitrocellulose, and probed with rabbit anti-DCM (PA5-144467), anti-Polr2b (PA5-21446), and anti-β-actin (CL594-60008) primary antibodies (1:1000, 1:1000, 1:2500) followed by HRP-conjugated anti-rabbit secondary (1:5000), with detection by enhanced chemiluminescence. DCM-Polr2b and endogenous Polr2b were resolved on the same membrane and distinguished by molecular weight ( 180 kDa and 150 kDa, respectively).

#### Doxycycline Calibration for efficient DCM methylation

For calibration experiments, dissociated NSCs at passage 5 were seeded into T25 flasks at a density of 2 × 10^5 cells/ mL in NSC proliferation medium. Cells were subsequently treated with Doxycycline (Dox) at final concentrations of 0, 1.3, 2, 4, 6, or 8 μg/mL for 12, 24, or 48 h. For *in vitro* chase experiments, cells were treated with the same Dox concentrations for either 12 or 24 h, followed by a 24- or 48-h chase period in Dox-free medium. Following treatment, cells were harvested and dissociated with Accutase to obtain a single-cell suspension. The GFP high fraction was then sorted by FACS into 384-well plates, with four minibulks per condition, for downstream analyses.

#### Coating for adherent assays

Plates were coated with poly-D-lysine (10 μg/mL; SigmaAldrich, P1024; overnight, room temperature), washed four times, air-dried, and coated with laminin (10 μg/ mL; Sigma-Aldrich, L2020; 2 h, 37 °C) immediately before seeding.

#### BMP4-induced quiescence, Doxycycline labeling, and reactivation

For adherent assays, NSCs were plated at 4 × 104 cells cm_−2_in complete medium. The next day, cells were washed once with PBS and switched to BMP4 medium [complete medium without EGF + 50 ng/mL recombinant human BMP4 (Bio-Techne)]. Doxycycline (4 μg/mL) was applied for 48 h starting 48 h after BMP4 exposure to induce DCM expression and DCM-dependent methyl labeling. After Doxycycline withdrawal, cells were either maintained in BMP4 medium or reactivated by removing BMP4 and switching to complete medium supplemented with 100 ng/ mL recombinant human Noggin (Bio-Techne).

#### Proliferating cohort labeling and delayed BMP4 exposure

In parallel, proliferating NSCs in complete medium received the same Doxycycline pulse (4 μg/mL, 48 h). After Doxycycline withdrawal, cells were either maintained in complete medium or transferred to BMP4 medium 48 h later.

#### Sample collection and FACS preparation

Cells were collected at TP0 (24 h), TP3 (96 h), and TP7 (8 days) after Doxycycline removal. Cells were dissociated with trypsin (5 min, 37 °C), washed twice with PBS, filtered through a 40-μm strainer, and resuspended in FACS buffer [PBS + 10% FBS] for sorting and downstream analyses. Sorting plates were preloaded with 3 µL lysis buffer per well, consisting of 2 µL RLT, 0.975 µL H_2_O, and 0.025 µL RNase inhibitor (40 U/µL).

#### FACS sorting of in vitro mouse NSCs

Cells were sorted using a BD S8 Discover equipped with a 100 µm nozzle. Live cells were identified by exclusion of Sytox Blue-positive events, and GFP-positive cells were selected to enrich *in vitro* cells with successful Doxycycline-induced reporter induction. Sorting was performed into 384-well plates, collecting both single cells and four minibulk samples per condition, with each minibulk consisting of 25 cells.

#### SVZ explant dissociation

SVZ explants were prepared from 2-month-old mice, including H2B-eGFP-DCM-Pol2B experimental animals as well as control animals used for GFP and non-GFP fluorescence-minus-one (FMO) gating. Explants were cultured ex vivo in explant medium consisting of 125 mL DMEM/ F-12, 125 mL Neurobasal medium, 2.5 mL GlutaMAX, 1.25 mL non-essential amino acids (NEAAs), 2.5 mL penicillin/ streptomycin, 2.5 mL N2 supplement, 5 mL B27 supplement, 87.5 µL β-mercaptoethanol (1:100 pre-diluted in DMEM), and 62.5 µL insulin. For culture, 1% Matrigel was freshly added to medium aliquots. Experimental explants were treated with Doxycycline at a final concentration of 8 µg/mL for 72 h. Explants were subsequently dissociated using the Neural Tissue Dissociation Kit (T) (Miltenyi Biotec, 130-093-231) and filtered through 40 µm strainers to obtain single-cell suspensions.

#### FACS sample preparation and staining

Cells were stained on ice for 20 min in the dark in the presence of Fc receptor block and washed twice with PBS containing 10% FCS. The antibody panel included CD9-Alexa Fluor 700 (Invitrogen, 56-0091-80), CD15-PE (Miltenyi Biotec, 130-114-011), O4-APC (Miltenyi Biotec, 130-118-978), PSA-NCAM-PE-Vio770 (Miltenyi Biotec, 130-095-212), CD45-APC-Cy7 (BD Pharmingen, 557659), and TER-119-APC/Cy7 (BioLegend, 116223). Sytox Blue was used for viability discrimination. Unstained, single-stained, and FMO controls, including GFP and non-GFP FMOs, were used to define gates.

#### FACS sorting of SVZ explant

Flow cytometric analysis and sorting were performed on a BD FACSDiscover S8. Viable singlets were gated after exclusion of CD45-and TER-119-positive events, and populations were assessed based on GFP, CD9, CD15, O4, and PSA-NCAM signal. Single cells were sorted into 384-well plates, and minibulk samples of 25 cells each were collected per marker. Sorting plates were preloaded with 3 µL lysis buffer per well, consisting of 2 µL RLT, 0.975 µL H_2_O, and 0.025 µL RNase inhibitor (40 U/µL).

#### Mouse pancreas isolation

Pancreatic tissue was collected from the above-mentioned mouse mice in parallel to the SVZ. The abdominal cavity was opened by a midline incision, the pancreas was exposed and carefully dissected, and excess surrounding adipose tissue was removed when necessary. Isolated tissue was kept on ice in PBS until further processing.

#### Mouse acinar-derived organoid and maintenance culture

The following sterile-filtered solutions were prepared in advance: solution S (4% bovine serum albumin [BSA] in PBS), solution R (1% BSA in PBS), solution D (1 mg/ml collagenase IV in 0.25% BSA in PBS), and solution W (2% penicillin-streptomycin in PBS).

Freshly isolated pancreatic tissue was transferred into 10 ml solution W and maintained on ice. Residual fat was removed, and the tissue was washed once again in 10 ml solution W. The pancreas was then minced into pieces of approximately 1 mm3. Tissue fragments were collected, washed in 10 ml solution W, and digested in solution D for 30 min at 37 °C in 5% CO2. During digestion, the suspension was triturated every 5 min using 5 ml serological pipettes to promote tissue dissociation.

The digested material was passed through a 100 µm cell strainer to deplete pancreatic islets, and the strainer was rinsed with 10 ml solution R. To enrich acinar clusters, four 15 ml centrifuge tubes were prepared with 6 ml solution S each, and 5 ml of filtered suspension was carefully layered on top of each to generate a BSA gradient. Samples were centrifuged once at 50 × g for 2 min at 4 °C. Supernatants were discarded, and pellets were sequentially resuspended and washed in solution S and then solution W, using the same centrifugation conditions (50 × g, 2 min, 4 °C).

After the final wash, each pellet was resuspended in 500 µl pancreatic organoid culture (POC) medium and pooled. POC medium consisted of DMEM/F12 + GlutaMAX mixed 1:1 and supplemented with 2% (v/v) B27 serum-free supplement, 1% (v/v) N2 supplement, 1% penicillin-streptomycin, 2 mM L-glutamine, 20 ng/ml recombinant human EGF, and 20 ng/ml recombinant human FGF2. At this stage, acinar cells remained in small clusters of approximately 4– 10 cells. Because preservation of cell-cell contacts supports organoid formation in this system, no additional dissociation was performed.

Acinar clusters were counted and combined with ice-cold, non-polymerized Matrigel at a final density of 250 cells/µl. For 3D culture, 20 µl droplets were plated into the appropriate culture vessels and allowed to polymerize for 20 min at 37 °C and 5% CO2 before addition of POC medium. Cultures were maintained at 37 °C and 5% CO2 in 24-well plates and treated for 72h with 8ug/ml Dox solution. The day of embedding in Matrigel was defined as day 0.

#### FACS sorting of mouse acinar-derived organoid cells

Flow cytometric analysis and sorting were performed on a BD FACSDiscover S8. Viable singlets were gated after exclusion of sytox blue-positive events, and populations were assessed based on size and GFP signal. Single cells were sorted into 384-well plates, and minibulk samples of 25 cells each were collected. Sorting plates were preloaded with 3 µL lysis buffer per well, consisting of 2 µL RLT, 0.975 µL H_2_O, and 0.025 µL RNase inhibitor (40 U/µL).

### Experimental work in zebrafish

#### Zebrafish husbandry

Zebrafish embryos and adults were kept according to the animal study protocol and research was performed in accordance with the ethical guidelines. The animal study protocol was approved by the Ethics Committee n°39 of Institut Pasteur (authorization #36936, April 26th, 2022) and DDPP-2021-021 of the Direction Départementale de la Protection des Populations de Paris. The following transgenic line was used: Tg(*sox2-2a-sfGFPstl84*) (*22*).

#### Zebrafish transgenesis

The “*pminiTol2_hsp70_DCM-Polr2bZebrafish-P2A-H2A-mCherry*” plasmid was generated by assembling the following five segments at equimolar concentration using the NEBuilder® HiFi DNA Assembly master mix (NEB #E2621) : (1) *pminiTol2* (gift from S. Ekker - Addgene plasmid #31829; http://n2t.net/addgene:31829; RRID:Addgene_31829[SO1.1]), linearized with NotI restriction enzyme (NEB #R0189S); (2) *hsp70l* promoter sequence, amplified by PCR from the plasmid “*p5E-hsp70l*” (Tol2kit #222[SO2.1]); (3) *DCM* sequence, amplified by PCR from the plasmid “*DCM-Polr2b_mouse*” (gift from J. Gribnau); (4) zebrafish *polr2b* sequence, amplified by PCR from the plasmid “*polr2b_zebrafish*”, obtained in our lab by cloning the *polr2b* sequence (RefSeq:NM_001024461) from zebrafish embryos cDNA using the following primers [forward: 5’-GGCGCGCCG-TATGATCAAGACGAAGATATCCAGTATG-3’; reverse: 5’-CAGGCTGAAGTTAGTAGCTCCGCTTCCCTTAAT-TAAGTCGCTGGTCATCATGCGCGG-3’]; (5) *H2A-mCherry* sequence amplified by PCR from the plasmid “*pME-H2amCherry[SO3*.*1]”*. All primers were designed using the NEBuilder® Assembly Tool. PCR amplifications were performed using the Phusion™ polymerase (Thermo Scientific #F530S). The *P2A* sequence was added into the primers connecting the last two fragments.

To generate the transgenic line, WT embryos were coinjected at one-cell stage with plasmid and *tol2* mRNA (30 pg mRNA and 20 pg DNA per embryo). F0 adults were out-crossed and their offspring heat shocked and subsequently screened for Cherry expression to identify founder fish. F1 embryos were raised to generate a stable transgenic line. Stable transgenic insertion is maintained as *Tg(hsp:DCM-polr2b-P2A-H2A-mCherry*).

#### *Genotyping of* Tg(hsp70:DCM-Polr2bZebrafish-H2A-mCherry)

Genotyping of adult fish was performed by PCR on tail samples. Before fin clipping, fish were anesthetized by immersion in system water containing 0.01% MS222 (Sigma-Aldrich, Cat# A5040). Fin clips were collected and DNA was extracted using the Phire Animal Tissue Direct PCR Kit (ThermoFisher scientific, Cat#F140WH) according to manufacturer’s instructions.Primers were chosen to bind inside the Cherry protein. The sequences for the primers are: Forward primer: 5’-GCTGAAGGTGAC-CAAGGGTG-3’; Reverse primer: 5’-GGCCTTGTAGGTG-GTCTTGAC-3’. An annealing temperature of 65C was used.

#### HCR and Immunohistochemistry

*DCM* mRNA expression was analyzed by in situ Hybridization Chain Reaction (HCR) RNA FISH version 3.0 (Choi et al, 2018). Fixation, detection and amplification were performed according to the HCR v3.0 protocol for whole-mount zebrafish larvae (Choi et al., 2016). After HCR, immunohistochemistry was performed against Cherry using Anti-RFP (abcam, ab6234) and incubated with 1:5000 DAPI overnight at 4°C. Embryos were mounted in Vectashield and imaged at LSM710 confocal microscope (Zeiss). Maximum intensity projections of z-stacks were generated in Imaris and representative images are shown in the figures.

#### Image acquisition

Confocal images for the HCR were acquired on a LSM710 confocal microscope (Zeiss) using a 40X oil objective (Plan-Apochromat 40×/1.3 Oil M27 – Effective NA between 1.3 and 1.4). Embryos were mounted in Vectashield. Images were acquired with a pixel size of 0.207 x 0.207 x 0.4 µm.

Overview images of the embryos in Fig. S3B,C were acquired on a fluorescent Stereomicroscope Olympus SZX16 equipped with a Olympus DP73 camera.

#### Image analysis of HCR

Images were processed with Bitplane Imaris software. Per embryo, one trunk region and one region in the retina were analysed. Per region, two substacks with a volume of 30 µm3 were created in Imaris. In these substacks, total intensity of the DCM HCR signal was measured using the surface tool. Sum intensity values of the *DCM* channel were then used for quantifications.

#### Induction of DCM labelling in zebrafish and collection of single-cells

To induce DCM expression, embryos were heat shocked if not otherwise mentioned with two pulses of each 45 min by replacing embryo medium with preheated embryo medium and then kept at 39C in an incubator. The first heat shock was started at 32 hpf and between the two heat shocks, the plates were transferred to 2C for 90 min. Embryos were screened for mCherry expression at a fluorescent stereomicroscope between 30 min - 2 hours after the heat shock.

#### Cell dissociation of zebrafish embryos

To obtain dissociated cells for FACS sorting, batches of 25 whole embryos at either 48 or 96 hpf were collected and euthanized with 4x tricaine on ice. Embryos were washed 2x with cold PBS and deyolked by triturating the embryos in 500µl deyolking buffer (55mM NaCl, 1.8mM KCl, 1,25mM NaHCO3 in H2O) until the yolk dissolved. Embryos were washed 2x in PBS. 96 pf embryos were then resuspended in 1ml TrypLE and incubated for 10 min on ice while gentry pipetted every 2-3 min. All embryos were centrifuged for 7 min, 300g at 4C, and then resuspended in 500 ul FACSmax, and incubated for 5 min at 30C. Afterwards the embryos were mechanically dissociated using 40µM cell strainers on ice. Upon centrifugation for 7 min, 300g, 4C, cells were resuspended in 600µl 1x dPBS with 0.04% BSA and kept on ice until FACS sorting.

#### FACS sorting of zebrafish cells

Cells were FACS sorted on a BD FACSAria™ III | High Sensitivity Flow Cytometer using a 70µM nozzle. To set the gates for mCherry and GFP expression, wild type embryos and embryos with the single fluorophore were used. 7-AAD was added to the cell suspension to exclude dead cells. At 48 hpf, all GFP+ cells were sorted, while at 96 hpf, only GFPlow cells were sorted to enrich the sort for mature neuronal cells. Single cells were sorted in a 384 well plate, in which we sorted 48 cells from 1 tube of 25 embryos.

### Preparation of libraries and scMT sequencing

*Library preparation for mouse cells and miniaturized scMT-seq protocol for profiling the transcriptome (cytoplasmic and nuclear mRNAs) and epigenome (DNA methylation and chromatin accessibility) of single cells*

We developed and implemented a miniaturized and higher throughput version of the scNMT-seq protocol (*20*). In this new version, the Smart-seq3 method and specific normalization steps were implemented. A detailed version of the protocol is described in (*20*). We used combinatorial indexing on the genomic DNA fraction, with a multiplexing capacity of 384 cells per run. The index combination and sequencing details are provided provided in (*20*)

#### Library preparation for zebrafish cells

We used the miniaturized scMT-seq protocol as described for mice, with the following adaptions: cycle numbers of cDNA amplification: 35; additional TSO reverse transcription as mentioned in the STAR protocol; amount of cDNA used for tagmentation:1 ng. We used combinatorial indexing on the genomic DNA fraction, with a multiplexing capacity of 384 cells per run. The index combination as used for the Mouse scMT.

The libraries were sequenced at a NextSeq2000 Sequencing System using the following kit: Sequencing kit RNA: P1 100 cycles - REF: 20074933 Illumina; Sequencing kit DNA: P3 200 cycles - REF: 20040560 Illumina.

### Computational Methods

#### RNA

##### Alignment

For transcriptomic reads we first extracted UMIs, using a custom based on code in the *umite* toolkit (*27*), and then aligned the reads to the mouse (GRCm39.107) or zebrafish (GRCz11.115) genome with STAR (v.2.7.0f; *28*). The counts were then collected with the umicount script of *27*. GFP and DCM sequences were added to the reference as separate chromosomes. The final count matrix was obtained by summing the unique intron and exon reads.

##### Clustering and visualization

The resulting count matrices were then processed with the Seurat (v.5.5.1; *29*) package. NormalizeData, FindVariableFeatures, ScaleData and RunPCA were run with the default parameters. For clustering we used the same neighbourhoods as for UMAP and applied the Louvain algorithm with a resolution chosen for each particular dataset. We kept only the cells with more than 5000 UMIs and, since we normally want to exclude spurious cells from further genomic sequencing, we also removed cells with an abnormally high distance to the nearest neighbour or an abnormally high ratio of the distances to the 1st nearest neighbour and to its 9th nearest neighbour thus identifying cells that are outside of its neighbour’s cluster). This filtering was not applied to the zebrafish dataset due to the presence of meaningful but lowly represented clusters. The exact threshold values, as well as parameters for the RunUMAP function and clustering, depended on the dataset and are provided in the corresponding scripts (see source code associated to this paper). Cluster annotation was performed manually based on known marker genes.

##### Differential Expression

In order to demonstrate the absence of a global effect of DCM methylation on the transcriptome (Fig. S4D), gene counts from the minibulk transcriptomes belonging to the same embryo pools were summed together, resulting in 2 biological replicates with and 2 without the DCM-Polr2b construct (all four replicates were heat-shocked). The resulting pseudobulks were then normalized with the estimateSizeFactorsForMatrix(type = “poscounts”) function of the DESeq2 package (v.1.50.2, *30*). The normalized expression was used to directly generate an MA-plot with

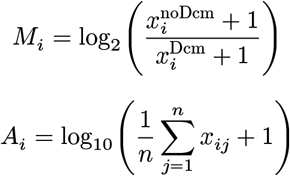

where *x*_*ij*_ is the normalized expression in gene *i* and cell *j*.

#### DNA

##### Alignment

Genomic data were first trimmed with Trim Galore (v.0.6.6; *31*). Biscuit (v.0.3.14; *32*) was used to align the reads, and we then used the provided bsstrand utility to identify the original methylated strand for each read, keeping only the reads for which this was possible. Since DCM methylation levels can be low in a cell, spurious calls may affect the signal; we therefore applied several filtering criteria at the per-read level:

- only reads with a mapping quality of 40 or higher were accepted;
- at least 70% of the read must be mapped;
- if a read contains at least 10 cytosines, no more than 70% of them may be methylated.

In addition, we excluded DCM motifs at the ends of a read and those whose middle base pair did not match the reference. The remaining motifs were collected, retaining the original methylation-strand information for the down-stream analysis. To aggregate BED files into per-region methylation counts, we used a script derived from an older version of MethSCAn (v.0.4.0; 33). Every chromosome was treated as two virtual chromosomes based on the methylation strand (e.g. instead of chromosome 1, the counts were assigned separately to chromosomes 1f and 1r).

For the wild-type dataset, additional filtering was applied to exclude DCM motifs that could potentially have been affected by GpC methylation.

##### Signal Strength

To estimate the cell’s DCM signal strength we use a Signal-To-Noise ratio (STN), which shows how much more strongly gene bodies are methylated compared to the “outside” regions. A DCM motif is considered “outside” if it is at least 5000 bp away from any gene, annotated promoter or enhancer (according to Ensembl annotation GRCm39.107). For the zebrafish dataset we do not have any information on the regulatory features, and therefore all motifs away from gene bodies were considered “outside”. First, we calculate the methylation fractions inside gene bodies and in the “outside” regions:

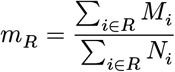

where *R* is the set of all DCM motifs in the region of interest (such as all gene bodies or all “outside” regions), *M*_*i*_ is 1 for a methylated motif and 0 for an unmethylated or unobserved one, and *N*_*i*_ is 1 for an observed motif and 0 for an unobserved one. The STN ratio is then calculated as the ratio of the gene-body methylation fraction to the outside methylation fraction.

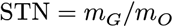

This ratio can be calculated for the entire cell or, for instance, for each chromosome separately, if required.

##### Strand splitting

Since DCM methylation marks are not copied to the newly synthesized strand, in a cell that has gone through at least one division cycle one may encounter drastically different methylation levels per strand. However, the sparse nature of WGBS data makes it unreasonable to split DCM methylation counts into two groups for each chromosome when there is no need to do so. Therefore, for each chromosome, we applied a two-proportion z-test to decide whether the global DCM methylation fractions on the forward and the reverse strands differ in a given cell.

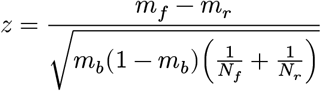

where *m*_*f*_ and *m*_*r*_ are the DCM methylation fractions on the forward and the reverse strands respectively, *m*_*b*_ is the methylation fraction on both strands combined, and *N*_*f*_ and *N*_*r*_ are the numbers of observed DCM motifs on the forward and the reverse strands respectively. From these z-values we calculated p-values and adjusted them for multiple testing using the Benjamini–Hochberg procedure. Only the chromosomes with FDR < 0.05 were considered different. For all other chromosomes the forward and reverse strands were pooled together for the downstream analysis. We excluded all the minor contigs, since their size makes the background estimate too noisy.

##### Deviance residuals

We assumed that any chromosome (and possibly any strand) in a given cell has a certain level of background methylation *p*_*bg*_, which is the probability for any DCM site to become methylated regardless of the transcription patterns in that cell. In practice we took the global methylation fraction (i.e. the one calculated over all the observed DCM motifs in the given chromosome/strand) as an approximation of this probability. Since background methylation levels vary drastically between cells, chromosomes and strands, we used deviance residuals as a background-independent measure of the gene’s methylation levels. If there is no transcription-related DCM methylation of a gene *j* in cell *i* located on chromosome *k*, we expect its methylation counts *M*_*ij*_ and *N*_*ij*_ (i.e. the total number of methylated DCM motifs in this gene and cell, and the total number of observed DCM motifs in this gene and cell respectively) to be binomially distributed with the background probability *p*_*ik*_

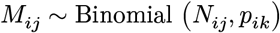

The deviance residuals then show how much our actual observations deviate from this distribution.

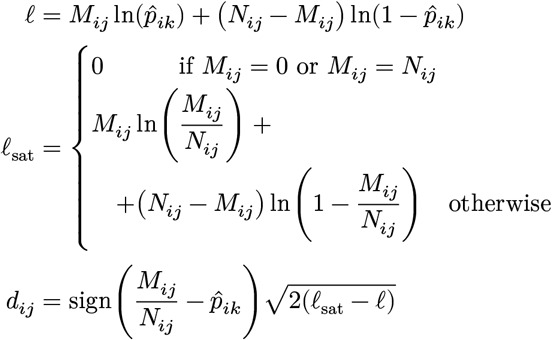

where 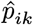 is the estimated background methylation probability (the global DCM methylation fraction). In cases where forward and reverse strands were considered to have different background methylation levels, for genes that have reads from both strands, deviance residuals were first calculated separately and then combined using Stouffer’s method.

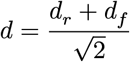

where *d*_*f*_ and *d*_*r*_ are the deviance residuals calculated for this gene based on the observations from the forward and reverse strands respectively. In the end we get no more than one value per gene and cell, whose absolute value is high if the gene is strongly methylated or undermethylated. These values now form our feature × cell matrix (with some values missing) that we can use for downstream analyses, such as dimensionality reduction or correlation with the known transcriptional signatures.

##### Feature filtering

It is especially important to remove “empty” chromosomes and/or strands from further consideration, since they not only increase noise but also generate incorrect signal by providing evidence that some genes are unmethylated. We therefore remove whole strands/chromosomes. To this end, we calculate a per-strand/chromosome STN ratio and keep only those where STN > 2 and at least 500 DCM motifs are observed, or where the global DCM methylation fraction is above 0.04. This is done because a low number of observed “outside” motifs may make the STN estimate noisy. These filtering criteria were applied only to the cells that show evidence of cycling (DCM methylation levels differ between strands in more than two chromosomes).

In addition, we exclude all genes where the total number of DCM sites (observed or not) is 5 or fewer, and genes that are observed in fewer than 30% of cells.

##### Cell filtering

Some filtering was also applied at the per-cell level.

Cells with fewer than 40,000 observed DCM motifs were considered to be of low quality and were therefore excluded entirely. These cells are not part of any plot or analysis in this paper. For the zebrafish dataset, the threshold was lowered to 10,000 observed motifs, due to the shorter genome and sparser DCM motifs. In the zebrafish dataset, one specific row (“O”) on the plate showed consistently and abnormally low DCM methylation levels. This row was excluded from any further consideration.

In order to avoid overloading dimensionality reduction approaches with cells without any detectable signal, we kept only cells where the promoter-based STN > 1.5. Promoters have very strong DCM methylation, and this threshold affects only extremely low DCM methylation levels. This filtering was applied only to the cells that do not show evidence of cycling (methylation levels differ between strands in no more than two chromosomes). Whenever this filtering is used in the paper, it is explicitly stated.

##### Iterative PCA

As one approach to reducing dimensions and subsequent visualization, we used the iterative PCA described in Ref (*33*). The matrix of deviance residuals was used as the input. The method starts by imputing zeros for all the missing values. At each step it calculates *n* principal components, then reconstructs the data based on them and updates the missing values according to the obtained reconstruction. The process stops either when the maximum number of iterations is reached, or when two consecutive reconstructions converge. For this project, we calculated 10 principal components.

#### Variational Autoencoder Motivation

As a more sensitive alternative to iterative PCA, we propose to use a shallow Variational Autoencoder. Its structure is based on our assumptions about the underlying process of DCM mark acquisition in a cell. We noticed that DCM methylation levels vary drastically per cell and that different methylation fractions (global methylation, gene-body methylation, “outside” methylation, etc.) are highly correlated with each other. This led us to assume that there exists an underlying “DCM activity” measure. Let us denote it as *b*_*i*_ for cell *i. c* is a coefficient that converts *b*_*i*_ into the number of times that the DCM-Polymerase construct approached the given DCM site due to background activity. *a*_*ij*_ defines the methylation signal of gene *j* in cell *i*, so that *a*_*ij*_*b*_*i*_ is the number of times that the DCM-Polymerase construct approached the DCM site while transcribing gene *j. d* is the probability for a DCM site to become methylated from a single approach of the DCM-Polymerase. From this we can estimate the probability of each DCM motif in gene *j*, cell *i*, not being methylated throughout the duration of the experiment.

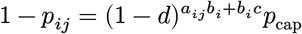

where 1 − *p*_cap_ is the probability of any DCM site in a cell being methylated (or being read as methylated) for any external reason, e.g. failed bisulfite conversion or CpH methylation (present in neuronal cells, etc.). This value “caps” our methylation probabilities from below, so we can call it the “capping” probability. We expect *p*_cap_ ≈ 1. Applying a complementary log-log transformation, we get:

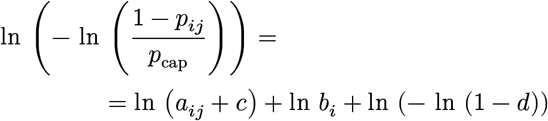

By merging the constants *c* and *d* with the signal and background coefficients respectively, we get:

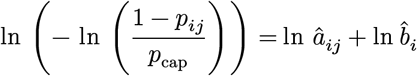

This formula provides a potential framework for extracting per-gene signal values. To this end we need to find the shared capping probability *p*_cap_, the background level *b*_*ik*_ (assuming an individual background per chromosome/ strand) and the signal coefficient *a*_*ij*_ that maximize the beta-binomial likelihood of our observations. Throughout this project we used *θ* = 40 as the concentration parameter of the beta-binomial distribution. The latent-space bottleneck of an autoencoder can prevent overfitting of these coefficients.

##### Simulated de-methylation

To help the autoencoder avoid picking up the DCM-activity signal (which is present in the data despite splitting the methylation input into signal and background components), we generate de-methylated copies of some cells. The autoencoder is then encouraged to place each copy close to its original. To ensure that the de-methylated levels resemble those present in the given dataset, we use “model” cells. First, we randomly assign a partner to each cell in the dataset. The one with the higher methylation becomes the “original”, and the less methylated one serves as the “model”. Therefore more highly methylated cells may have multiple copies, while some less methylated ones may get no copies at all. But since we cannot simulate methylation acquisition without distorting the unknown true signal, we have to accept this asymmetry. The following steps are performed per gene. Next, we modify the per-strand distribution of the original’s counts to match the model. If the model has the strands pooled for some chromosome, we pool the original’s counts as well. If the model has the strands split and the original does not, we randomly split the original’s counts between the two strands by separately splitting the methylated and unmethylated motifs.

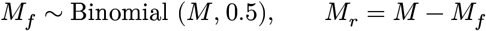

where *M* is the original’s number of methylated motifs, and *M*_*f*_ and *M*_*r*_ are the simulated numbers of methylated motifs on the forward and the reverse strand respectively.

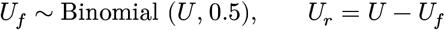

where *U* is the number of unmethylated DCM motifs. Then the number of observed motifs is reconstructed as:

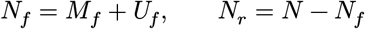

Now the actual demethylation occurs. We assume that the model’s chromosome/strand methylation is lower than the original’s. If this is not true, we leave this chromosome/ strand unaltered. The demethylation must respect the capping probability *p*_*cap*_, since this is required for the copy’s STN to approximate that of the model. We can decompose the fitted probabilities as *p*_*ij*_ = (1 − *p*_cap_) + *p*_cap_*Q*_*ij*_, where *Q*_*ij*_ is the DCM-dependent part. It depends on 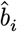 1and is therefore expected to be lowered by the demethylation. If *λ* is the fraction of the signal that remains, the methylation fraction of a feature becomes

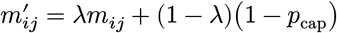

We can then obtain the new number of DCM methylation marks by drawing from a binomial distribution independently for each feature

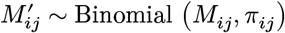

where *π*_*ij*_ is calculated as
}

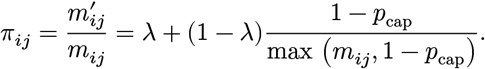

Features at or below the floor 1 − *p*_cap_ are left intact. *λ* is set per chromosome/strand so that the copy reaches the methylation level *m*_*m*_ of its model cell, which means 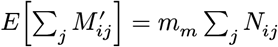. This leads to

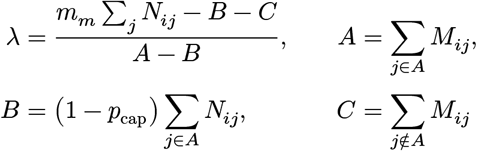

where *A* is the set of all features with methylation above the fixed floor. *λ* is then clipped to [0, 1].

##### Background fit

Before training the autoencoder, we first fit the background parameters *p*_*cap*_ and *b*_*ik*_. To this end we assume that there is a single gene coefficient *a*_*j*_ for each gene, shared across all cells, and find the optimal combination of all the parameters (the one that maximizes the betabinomial likelihood). In order to obtain more general background parameters, we perform a nested optimization. For the current values of the background parameters, we roughly optimize the gene coefficients (with a small number of steps). We then use these gene coefficients to take the next optimization step for the background parameters. After this process the background parameters are fixed, and will be used in the autoencoder’s training as external parameters for the decoder, the fitted gene parameters will be used as a fixed baseline during the reconstruction

Since the value of capping probability is required for the downmethylation procedure, we run one extra background fitting round as the first step. Then, after the simulated cells are added to the dataset, the background fit is run again.

##### Encoder

The encoder consists of standard sequential blocks of Linear, BatchNorm (optional, not used for the final runs in this project), ReLU and Dropout layers, each block reducing the number of features to a preset value (we used 128d and 32d blocks). Let *M*_*ij*_ and *N*_*ij*_ denote the number of methylated and observed DCM motifs in cell *i* and gene *j*, and let *R*_*ij*_ = *M*_*ij*_/(*N*_*ij*_ + *ϵ*) be the corresponding methylation rate (*ϵ* = 10^−8^). Genes on different strands are treated as independent features at this stage, so each gene *j* contributes up to three features (forward, reverse, and both), some of which may be unobserved (*N*_*ij*_ = 0). Rather than passing the raw rates to the encoder, each feature is replaced by its coverage-shrunk deviation from the background methylation level of its own chromosome/ strand. The signal value for the encoder is then calculated as:

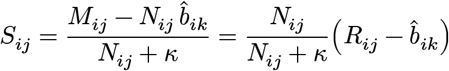

where 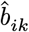 is the strand’s background, estimated as the DCM methylation fraction across all the features on the given strand, and *κ* is a shrinkage-strength parameter (*κ* = 2). In this way the encoder receives some information on the background and is encouraged to rely more on genes with a higher number of observations, which are considerably less noisy. Unlike in the case of raw rates as input, 0 becomes a consistently neutral value that indicates the lack of evidence of extra (or missing) methylation. In order to further discourage the encoder from learning from genes with a low number of observations, which can misshape the latent space during the early epochs and leave it stuck in this configuration, we apply an extra observation-based dropout procedure. During training, each feature can be randomly dropped out with probability:

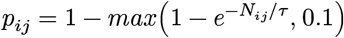

where *τ* is an external parameter (*τ* = 10). Finally, to reduce the number of input dimensions, the signal from all three strands (forward, reverse and both) is pooled together:

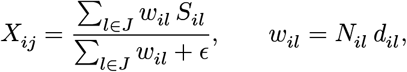

where *J* is the set of all strands for gene *j*, and *d*_*il*_ indicates whether the given feature is kept for the current training epoch. Although the two steps — background correction and strand pooling — are implemented as separate features that can be turned on and off individually, background correction is required for strand pooling, since the neutrality of 0 is crucial for this step.

##### Latent space

In the latent space all points are forced onto a *D*-dimensional unit hypersphere, where *D* is the preset number of latent-space dimensions (*D* = 10). To add a variational component to the autoencoder, we assume a spherical Cauchy distribution and perform a sampling step by drawing from a uniform distribution and applying a Möbius transformation (the approach was proposed in *34*).

##### Decoder

The decoder has the same structure as the encoder, but in the reverse order. It is tasked with decoding the latent-space coordinates into a set of gene coefficients *a*_*ij*_, with the strand information no longer taken into account.Therefore, if strand pooling is not applied, the decoder is three times smaller than the encoder. Otherwise they are identically structured.

##### Loss functions and training

To train the autoencoder we use three loss functions. The reconstruction loss is calculated as the negative log beta-binomial likelihood, with the estimated probabilities calculated according to formula (x). It uses the *a*_*ij*_ coefficients as reconstructed by the decoder, together with the prefitted background parameters *p*_cap_ and *b*_*ik*_. The demethylation loss is the cosine distance between the positions of the original cells and their demethylated copies in the latent space. We use their deterministic mean positions obtained from the encoder, rather than the resampled ones. For this loss the originals’ gradients are detached to ensure the copy is pulled towards its original. The KL loss is calculated according to (*34*). It depends on the fitted mean positions of each cell and its concentration *ρ*, and pulls the combined posterior towards the uniform distribution on the hypersphere. The gene coefficient penalty is the L2 norm of the difference between the fitted *a*_*ij*_ coefficients and the baseline obtained during the background fitting. This ensures that genes with a low number of observations stay close to the baseline instead of taking strongly varying and poorly constrained values.

The training process is split into three phases. During the first phase the reconstruction is calculated only from a set of around 2500 genes with the highest deviance. Generally, the best cluster separation is obtained during this step. In fact, if that is the goal, we recommend using only a limited set of important features. However, we have noticed that such a feature set, defined based on the methylation values alone, often has a small intersection with the variable features based on the transcriptome. Therefore, a second phase is added to the autoencoder. During this phase all other features are gradually opened up to the reconstruction loss. In order to protect the already obtained structure of the latent space from the influx of noise, another anchor penalty is added during this phase. It is the cosine distance of the cells from their positions at the end of the first phase. The final third phase is training with a frozen encoder, and it serves to refine the fitted gene coefficients.

##### Outputs

As the outputs of the trained autoencoder, we use the following values:

- the latent space (deterministic, not resampled) is used as the reduced-dimension space for further visualization or clustering.
- the matrix of fitted gene coefficients for all the cells is used to identify cluster-specific markers or correlations with known transcriptomes.
- the spherical Cauchy concentration is used to identify cells that the model failed to fit properly due to a lack of signal; we use the cut-off around 0.5, which can be visually identified by plotting the concentration vs. the global DCM methylation fraction or STN ratio of all the cells.

The exact run parameters can be found in the provided notebooks.

#### CpG Methylation

CpG methylation was processed by running the standard Biscuit pipeline (*32*). The pileup and vcf2bed utilities were used to obtain BED files with all the CpG site counts per cell. After that, the standard MethSCAn pipeline (*33*) followed: CpG counts were stored as matrices, variably methylated regions were detected, and then shrunken residuals were calculated for the detected regions. The dimensionality of the shrunken-residual matrix was reduced with iterative PCA, and the resulting 10-dimensional space was visualized with UMAP.

## Notes

### Competing Interest Statement

The authors have declared no competing interest.

