## Supplementary material for "Joint recording of past and present transcriptomes reconstructs single-cell trajectories": Full supplementary material

### SUPPLEMENTARY TEXT

#### The zebrafish and mouse genomes are suitable for the recording of DCM marks in single cells

DCM recognizes and methylates the palindromic motif 5'-CC[A/T]GG-3', independently on both strands. This motif appears on average every 465 bp in the zebrafish genome and every 173 bp in the mouse genome. DCM motifs appear in clusters (Fig. S1A,B), as previously suggested (*19*), but there is no bias in the number of DCM motifs per chromosome when normalized by chromosome length (Fig. S1C,D). The proportion of DCM motifs located inside genes vs in intergenic regions reflects the relative sizes of these domains in the genomes, showing no specific enrichment or depletion of DCM motifs inside genes (Fig. S1E,F). Finally, the distribution of DCM motifs per gene displays a broad range of values (Fig. S1G,H) but is proportional to gene length (Fig. S1I,J). In genes, we also observed a notable enrichment for DCM sites located in the coding vs 5' and 3' UTR sequences (Fig. S1E,F). Finally, using EnrichGO, we ruled out that genes with especially low DCM labeling potential (notably due to their small size) could belong to specific biological functions (Fig. S1K-N).

Thus, the DCM labeling potential of mouse and zebrafish are qualitatively similar, and both are suitable to exploit DCM labeling of active genes expressed in all biological processes and tissues, including the neuronal lineage, of particular interest here.

#### Use of scMT-seq

We initially used the scNMT-seq protocol (*20*), which comprises sequencing of mRNA (transcriptome, "T") and bilsufite-converted DNA (cytosine methylation, "M"), preceded by application of a GpC methyltransferase to label accessible chromatin (nucleosome positions, "N"). However, up to 10% of DCM motifs in wild-type cells without DCM were methylated due to off-target GpC methyltransferase activity. As a consequence, we omitted the "N" step (scMT-seq) in the present study.

### SUPPLEMENTARY FIGURES

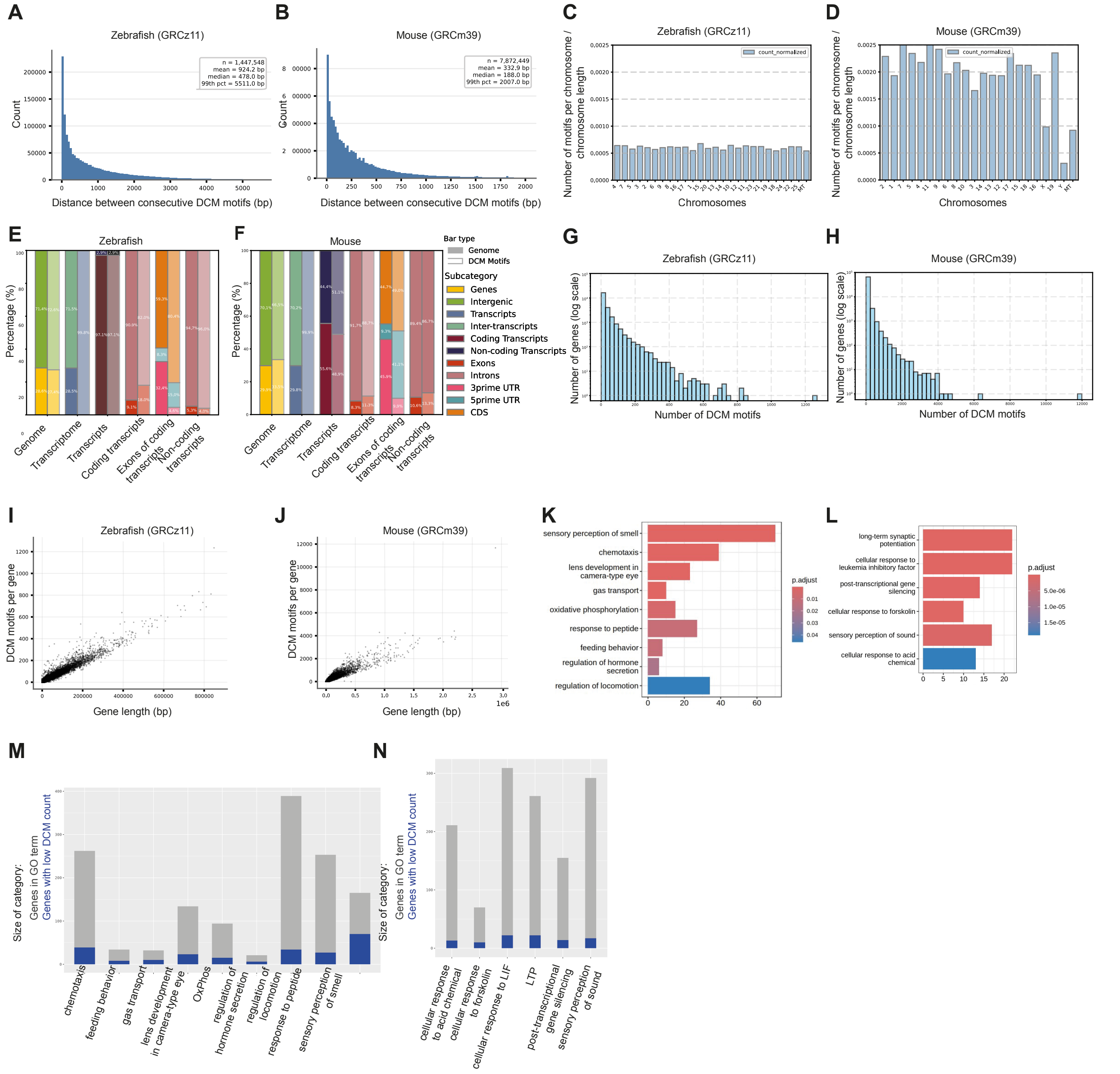


**Supplementary Figure S1. Genome analysis of DCM motifs in zebrafish and mouse. (A-B)** Distribution of inter-motif DCM distances in the zebrafish (A) and mouse (B) genome. **(C-D)** DCM motif density per chromosome normalized to chromosome length in the zebrafish (C) and mouse (D) genome**. (E-F)** Proportion of genomic categories (filled bars) versus DCM motif distribution across these categories (hatched bars) in the zebrafish (E) and mouse (F) genome. **(G-H)** Distribution of DCM motif number per gene in zebrafish (G) and mouse (H). **(I-J)** Number of DCM motifs in genes is proportional to gene length in both zebrafish (I) and mouse (J). **(K-L)** GO term enrichment analysis of genes with a low number of DCM motifs (a number inferior to the 0.05 quantile of DCM occurrence per gene) in the zebrafish (K) and mouse (L) genome. **(M-N)** Proportion of low-DCM-motif genes per enriched per GO term in the zebrafish (M) and mouse (N) genome.

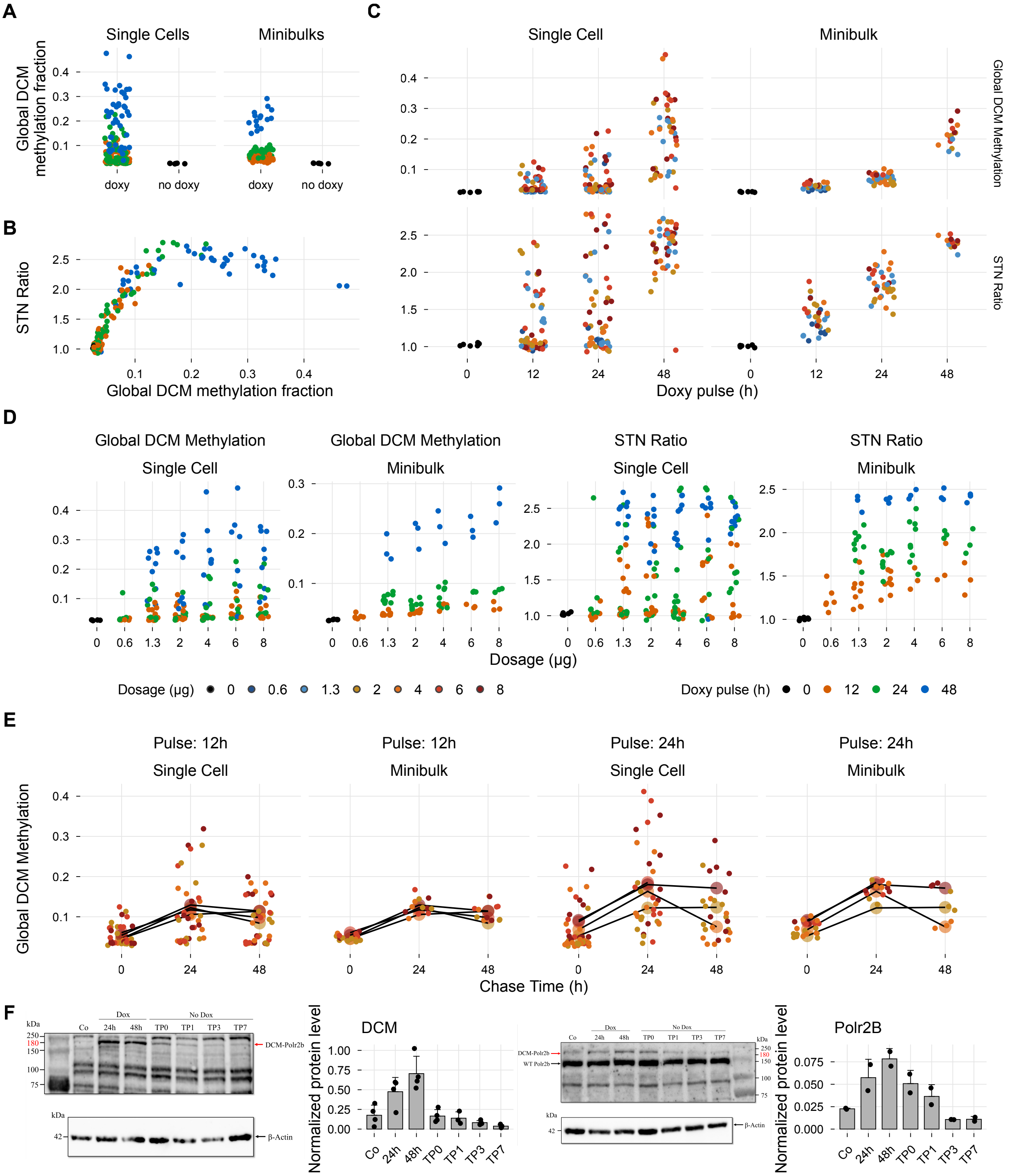


**Supplementary Figure S2. Optimization and temporal dynamics of DCM methylation labeling in mouse. (A)** Global DCM methylation levels following doxycycline (Dox) pulse labeling in mice. **(B)** Signal-to-noise ratio (STN) of DCM methylation. **(C)** Effect of Dox pulse duration on STN and global DCM methylation. Both increased markedly with pulse duration. Bulk measurements showed lower variability and provided population-level estimates, whereas single-cell measurements revealed cell-to-cell heterogeneity. **(D)** Dose dependence of STN and global DCM methylation. Doses >1.3 µg produced only modest additional increases, most apparent at shorter pulse durations. **(E)** Temporal dynamics of global DCM methylation following Dox withdrawal. Methylation continued to increase during the first 24 h after withdrawal and declined after 48 h, indicating that acquisition of new DCM methylation largely ceases 24 h after the Dox pulse. **(F)** Western blot analysis of DCM-Polr2b and endogenous Polr2b in DCM-Polr2b NSCs treated with Dox (4 µg/mL, 48 h). Lysates were collected after 24 and 48 h of treatment and at 0, 1, 3, and 7 days after Dox withdrawal (Tp0, c1, c3, and c7), with untreated cells as negative control. DCM-Polr2b ( 180 kDa, red arrow) and endogenous Polr2b ( 150 kDa, black arrow) are indicated; β-actin (42 kDa) served as loading control. Molecular weight markers (kDa) are shown at right. Quantification is shown in the right panel.

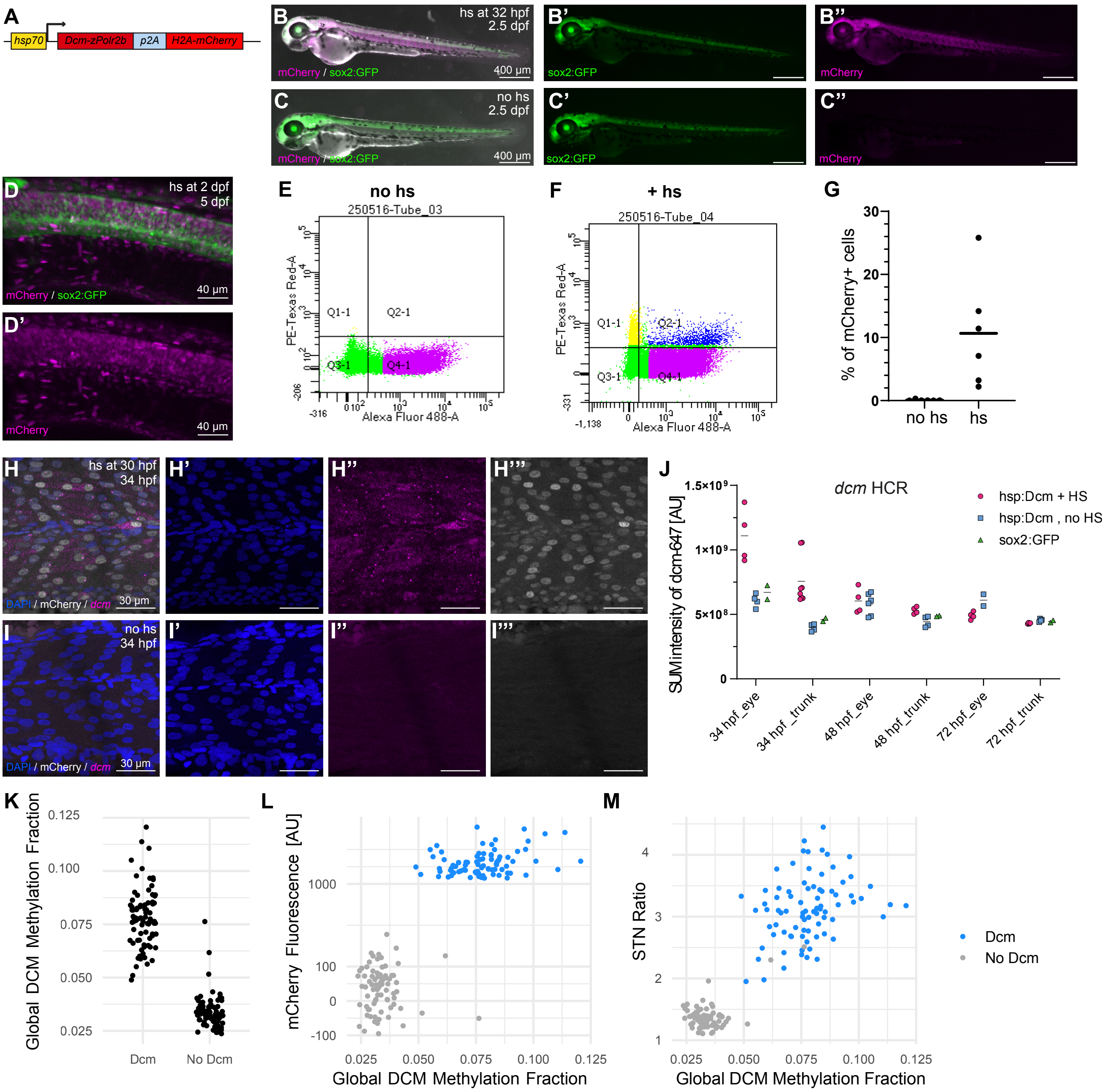


**Supplementary Figure S3. Characterization of the heat-shock-inducible DCM transgenic reporter line in zebrafish. (A)** Schematic of the genetic construct of Tg(*hsp:DCM-polr2b-P2A-H2A-mCherry*) showing the heat-shock-inducible promoter (*hsp70*), the DCM–POLR2B fusion (bacterial DNA cytosine methyltransferase fused to zebrafish RNA Polymerase II subunit B), a self-cleaving P2A peptide, and a histone H2A–mCherry nuclear reporter. **(B-C)** Stereomicroscope images of mCherry expression in Tg(*hsp:DCM-polr2b-P2A-H2A-mCherry*) representative embryos upon heat shock (B), compared to non-heat-shocked controls (C) at 2.5 dpf. (N=2, n =4). **(D)** Confocal image of a trunk region of a representative embryo at 5 dpf following heat shock at 2 dpf. Scale bar = 400 µm. (N=2, n =4). **(E-F)** FACS plot showing mCherry expression of Tg(*hsp:DCM-polr2b-P2A-H2A-mCherry*) cells at 48 hpf with (E) and without (F) heat shock induction at 32 hpf. mCherry+ gate was defined based on wild type controls. **(G)** Quantification of mCherry+ proportion among sorted cells. **(H-I)** HCR for *dcm* mRNA in 34 hpf Tg(hsp:DCM-polr2b-P2A-H2A-mCherry) embryos heat-shocked at 30 hpf (H) compared to non-heat-shocked controls (I) (N=2, n ≥2). Different patterns of mRNA expression reflecting the nascent transcript in the nucleus versus mature transcripts in the cytoplasm correlating with $\text{mCherry}^{\text{high}}$ cells can be observed. Scale bar = 30 µm. **(J)** Quantification of mean *dcm* HCR fluorescence intensity at 34, 48 and 72 hpf in heat-shocked vs. control embryos. No increase in not induced transgenic animals compared to controls is visible. Bars represent mean; (N=2, n ≥2). **(K)** Global DCM methylation level upon heat shock induction in zebrafish. **(L)** Scatter plot showing the correlation between DCM methylation fraction and mCherry intensity recorded by flow cytometry in the same cells at 48 hpf. Cells are colored by DCM-induced (+DCM) or control (–DCM) status. **(M)** Signal-to-noise ratio of DCM methylation at 48 hpf.

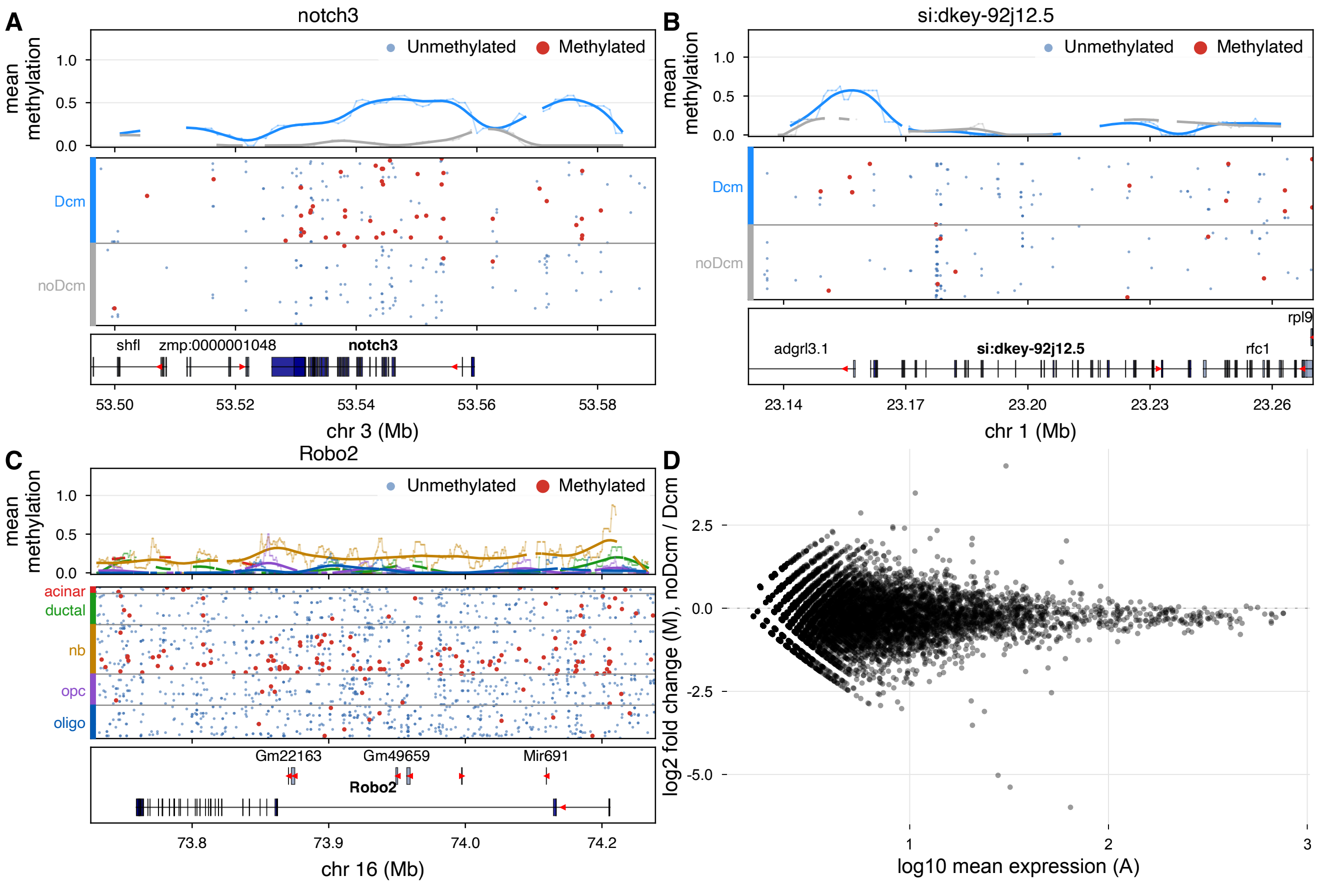


**Supplementary Figure S4. DCM labels specifically actively transcribed genes. (A)** Genome browser view of DCM methylation reads in zebrafish *notch3* gene. **(B)** Genome browser view of DCM methylation reads in zebrafish *si:dkey-92j12.5* gene, expressed in photoreceptors. **(C)** Genome browser view of DCM methylation reads in mouse *Robo2*. **(D)** RNA-seq analysis comparing average gene expression values before and after induction of DCM expression in zebrafish.

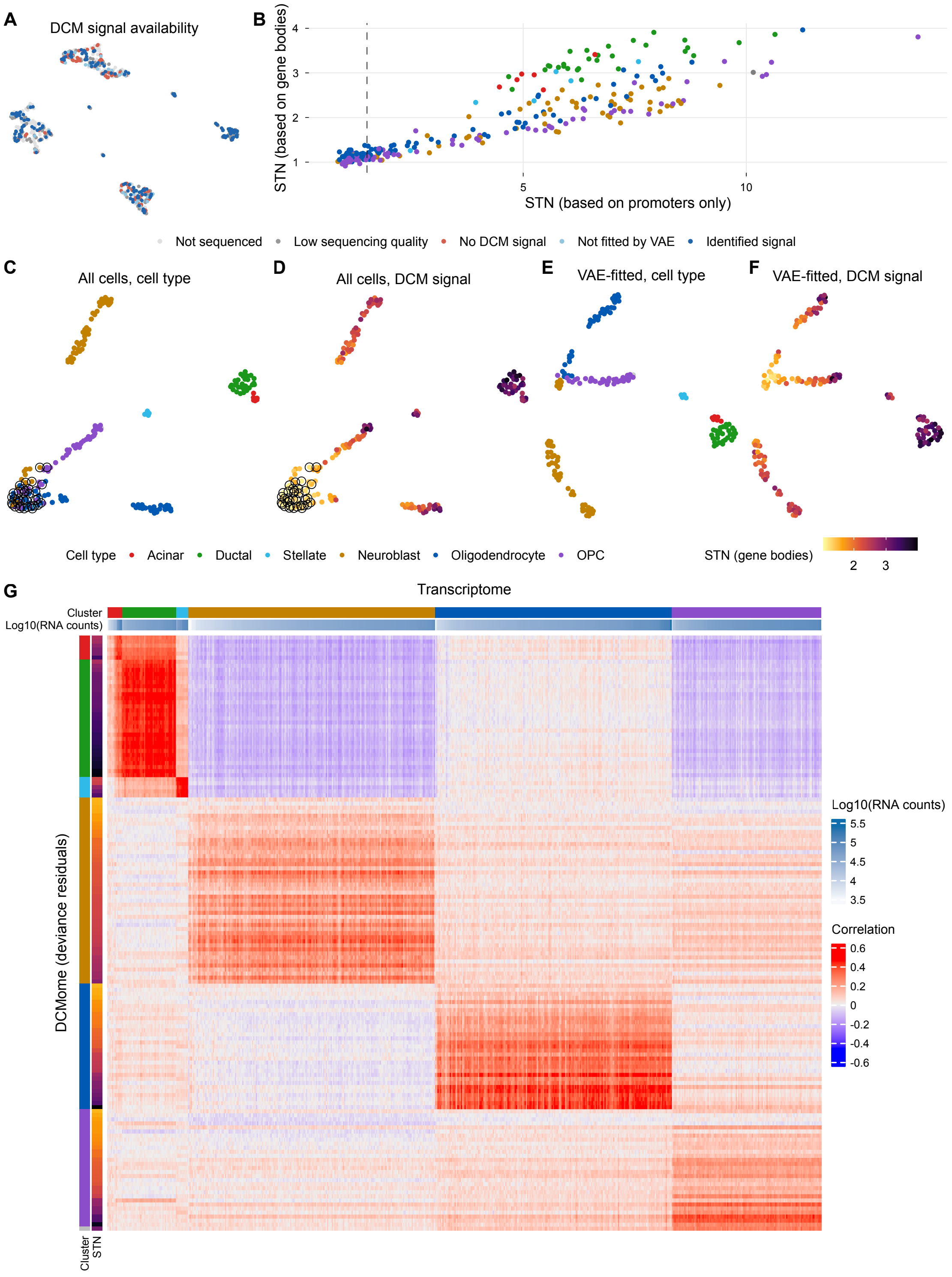


**Supplementary Figure S5. scDCMome quality control, representation, and cell-state mapping in mouse pancreatic and brain cells in vitro. (A)** DCMome signal quality for cells shown in Fig. 1C, displayed on the transcriptome-based UMAP from Fig. 1C. Cells are colored according to their suitability for downstream analyses (Fig. 1D,E). Grey indicates cells that were not sequenced or for which sequencing was unsuccessful, orange indicates cells without detectable DCM signal, and light blue indicates cells excluded because of low signal. **(B)** Gene body-based versus promoter-based signal-to-noise (STN) ratios for ADO and SVZ cells. Promoter methylation showed higher signal and specificity than gene body methylation and was therefore used as an early indicator of DCM activity. Cells below the indicated threshold (STN_prom $< 1.5$; dashed line; $n = 78$) were excluded from downstream analyses. **(C,D)** UMAP representation of the latent space obtained by PCA of binomial deviances derived from DCM methylation counts for all cells shown in Fig. 1D, colored by transcriptome-defined cell type (C; Fig. 1C) or STN ratio (D). Red-circled cells correspond to low-signal cells excluded from Fig. 1D and shown in light blue in (A). **(E,F)** UMAP representation as in (C), restricted to cells retained for the analysis in Fig. 1D, colored by transcriptome-defined cell type (E) or STN ratio (F). **(G)** Analysis as in Fig. 1E using raw binomial deviance residuals rather than VAE-denoised DCMome vectors. Correlations are reduced in the absence of denoising.

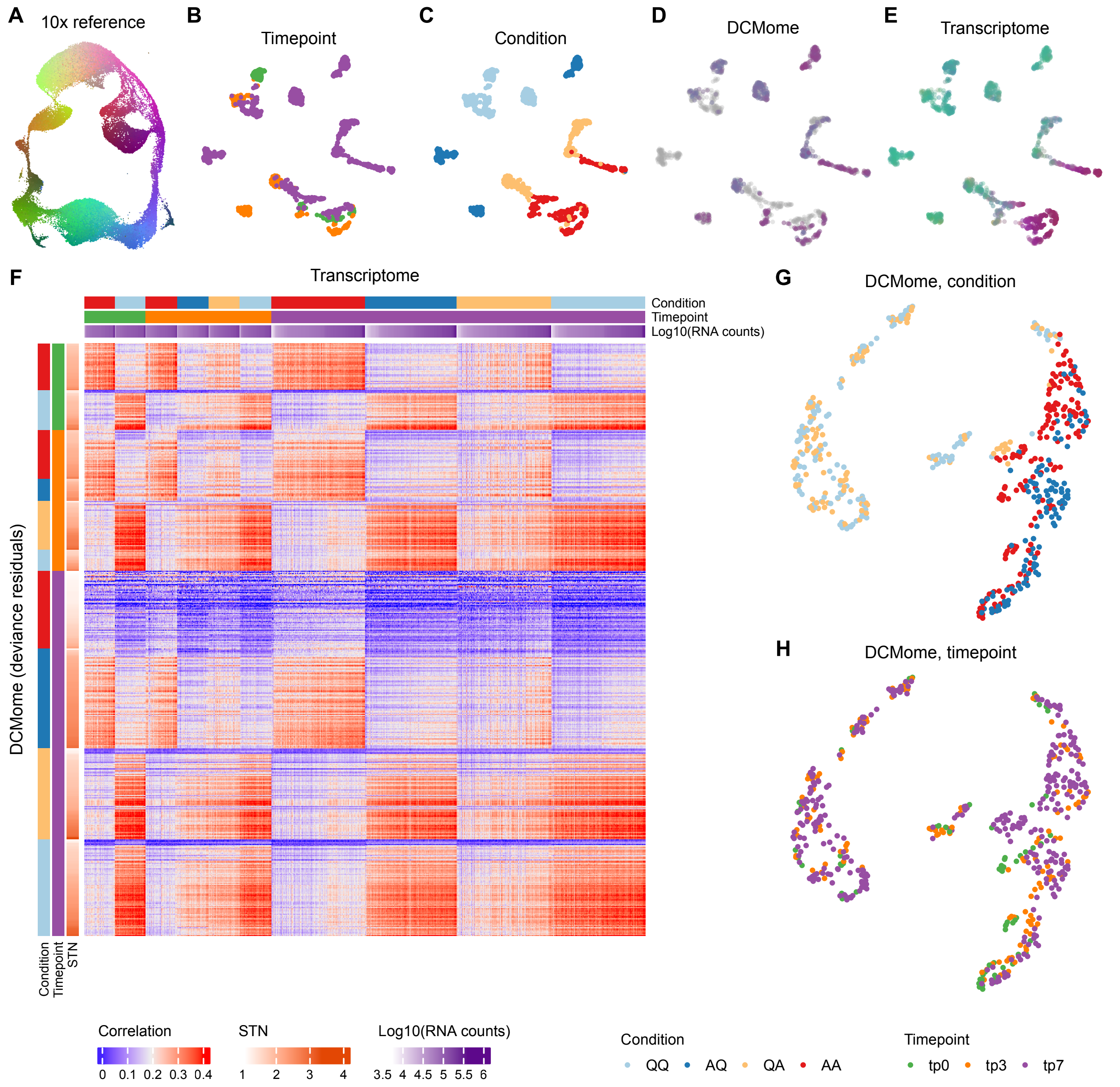


**Supplementary Figure S6. Mapping of mouse NSC culture pulse-chase cells onto the 10X SVZ atlas. (A)** SVZ atlas UMAP as in Fig. 2F, with each atlas cell assigned a unique color. **(B,C)** UMAP of cultured NSCs colored by time point (B) and condition (C). **(D)** Same as in (B), with atlas mapping based on the highest DCMome correlation. **(E)** Same as in (B), with each cell colored according to the atlas cell showing the highest transcriptomic correlation. **(F-H)** The VAE-based analysis can compensate for signal loss due to cell divisions. (F) As Fig. 2E, but with binomial-deviance/PCA DCMome profiles instead of VAE outputs. (G,H) As Fig. 2C,D, but using a binomial-deviance/PCA latent embedding instead of VAE.

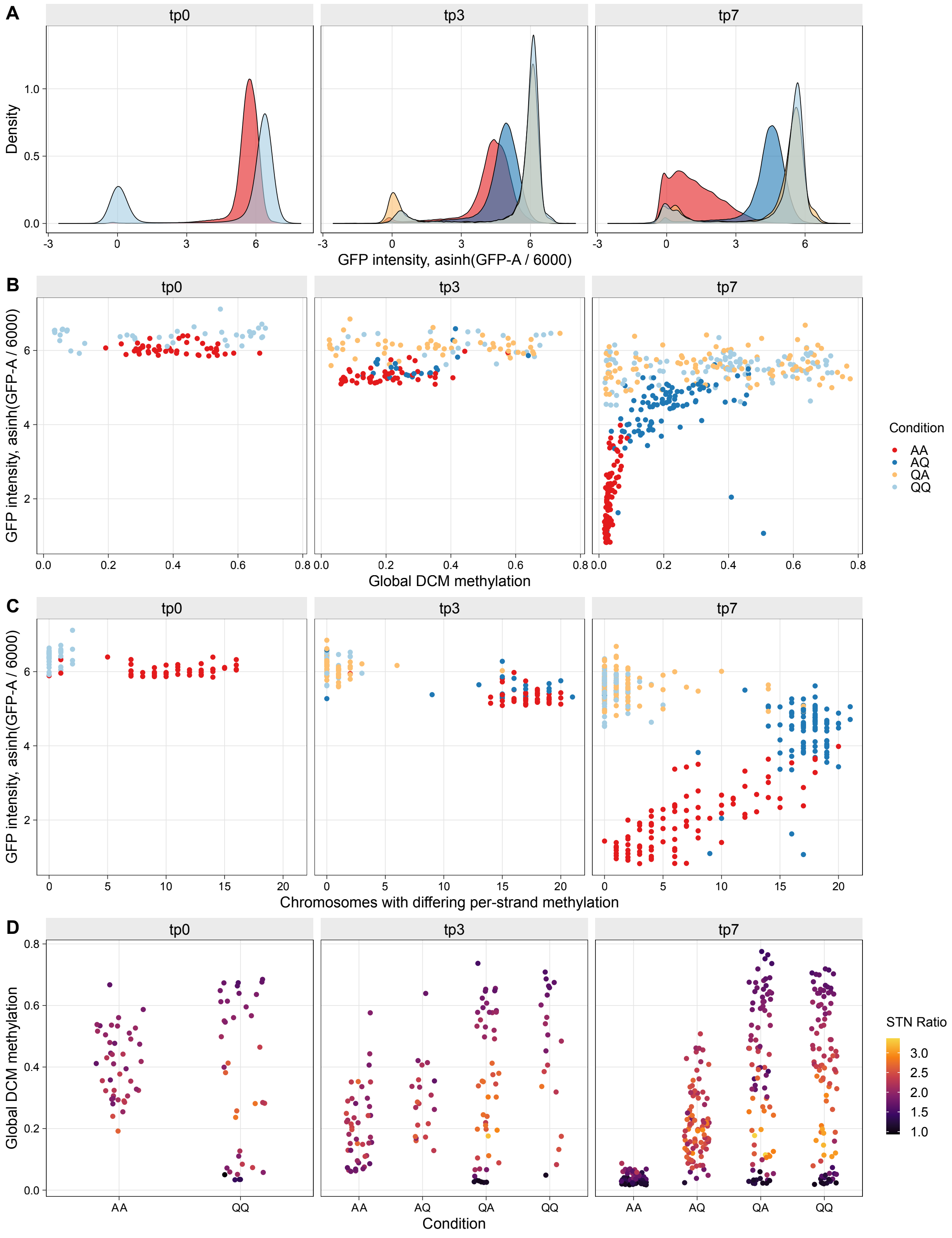


**Supplementary Figure S7. Changes in GFP emission and DCM methylation across cell divisions in mouse NSCs in vitro. (A)** Density plots of GFP emission for QQ and AA cells at t0, and QQ, AA, QA, and AQ cells at tp3 and tp7. GFP declines more sharply in AA/AQ than in QA/QQ, which largely overlap. **(B)** GFP intensity vs. DNA methylation (β value) at tp0, tp3, and tp7. By tp3, AA/AQ reach 50% of the β values seen in QQ/QA; by tp7, AA remains low while QA approaches QQ levels. **(C)** Number of chromosomes with strand-discordant DCM methylation (arising during the cell cycle, when newly synthesized strands lack DCM marks). Cells with high GFP have few discordant strands. As cells cycle, GFP declines and discordant-strand counts rise, until both strands lose DCM entirely, at which point GFP continues to decline while discordant-strand counts fall back down. **(D)** Distribution of global DCM methylation and DCM signal-to-noise (STN) ratio, stratified by sample, as in Fig. 2B.

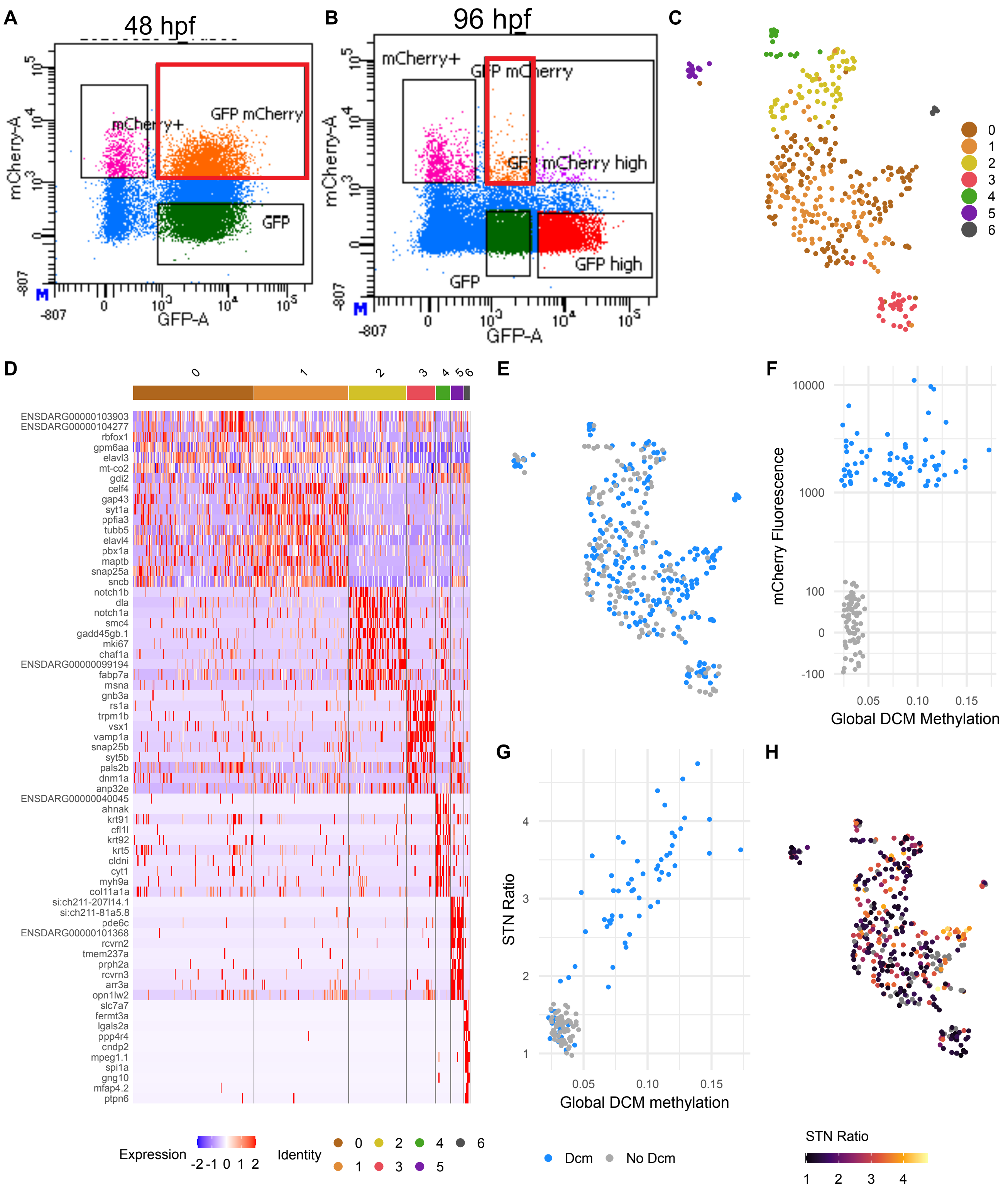


**Supplementary Figure S8. Technical validations in zebrafish.** **(A-B)** FACS gating strategy for GFP-expressing cells. (A) At 48 hpf, all GFP+ cells were sorted; (B) at 96 hpf, only GFP$^{low}$ cells were included (red box). **(C)** Transcriptomic UMAP of sorted single cells from 48 hpf and 96 hpf, colored by unsupervised cluster identity. **(D)** Heatmap showing the top 10 differentially expressed genes (ranked by adjusted p-value) for each unsupervised cluster shown in (C). **(E)** Transcriptomic UMAP colored by DCM-induced cells (Dcm) and control cells without DCM induction (noDcm). **(F)** Scatter plot showing the correlation between DCM methylation fraction and mCherry intensity recorded during FACS of 96 hpf cells. Cells are colored by DCM-induced (Dcm) or control (noDcm) status. **(G)** Signal-to noise ratio of cells at 96 hpf. **(H)** Transcriptomic UMAP of cells at 48 hpf and 96 hpf colored by DCM signal-to-noise ratio.

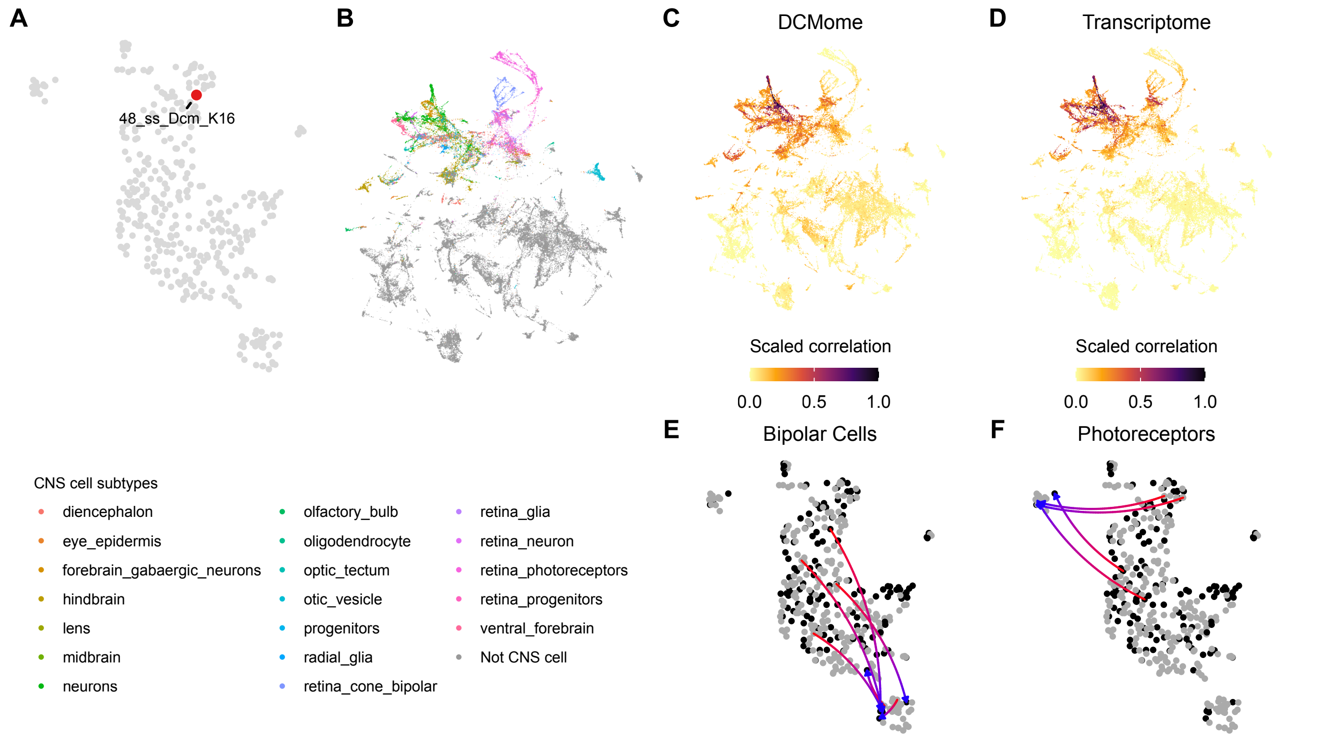


**Supplementary Figure S9. Examples showing showing the identificatin of cell types and the prediction of the state of origin by the DCMome in zebrafidh in vivo. (A-D)** Position of a single progenitor cell (48_ss_DCM_K16) highlighted in the UMAP with a red dot (A). Correlation of the cell's DCMome (C) and transcriptome (D) to transcriptomes of Zebrahub atlas cells (B) is shown as a heatmap projected onto the Zebrahub UMAP, where darker colors indicate higher correlation. CNS clusters are shown in color; all other cell types are shown in grey (B). **(E,F)** UMAP as in Fig. 4H,I but where grey indicates cells profiled for the transcriptome only, and black indicates cells profiled for both the transcriptome and DCMome. Arrows connect the transcriptomic position of a given cell (blue arrowhead) to the cell in the UMAP whose transcriptome most closely correlates with that cell's DCMome (red arrow base). Arrows are shown for: bipolar cells (E) and photoreceptors (F).
